# Spatial and Single-Cell Dissection of Fibroblast Subpopulation Reprogramming Driving Stromal Collapse in Breast Cancer Lymph Node Metastasis

**DOI:** 10.1101/2025.08.18.670793

**Authors:** Hadith Rastad, Mahnaz Seifi Alan, Mahin Seifi Alan, Sumeet pandey

**Author notes:** corresponding authors, Cardiovascular Research Center, Alborz University of Medical Sciences, Karaj, Iran; &; Postal code: 3149779453. Mahin Seifi Alan and Hadith Rastad contributed equally.

## Abstract

**Objective:** This study aimed to define conserved molecular drivers of breast cancer lymph node metastasis (LNM) through an unbiased, multi-omics approach, resolving spatial and cellular mechanisms of stromal reprogramming.

**Methods:** We integrated three bulk RNA-seq datasets (GSE110590, GSE193103, GSE209998) to identify differentially expressed genes (DEGs) in LNMs versus primary tumors. Tissue-specific genes were excluded where possible. Single-cell (scRNA-seq; GSE225600) and spatial transcriptomics (Visium) from primary/metastatic tissues were analyzed using Seurat and spacexr. Fibroblast subclusters were isolated for functional enrichment. Clinical validation included genetic alteration profiling (cBioPortal), survival analysis (KM Plotter), and stage-specific expression dynamics (TNMplot). Therapeutic candidates were screened via CTD.

**Findings:** Bulk RNA-seq identified three genes consistently downregulated in lymph node metastases compared to primary tumors: MAB21L1, F2RL2, and COL6A6 (log_2_FC ≤ −2.62; adj. p < 0.05). Single-cell RNA-seq analysis revealed that all three DEGs converged exclusively within specific fibroblast subpopulations in primary tumors; these DEGs were not enriched in other cell types. DEG-expressing fibroblasts orchestrated complementary protective barriers: F2RL2^+^ fibroblasts mediated immune crosstalk (via CXCL14, C1S) and TGF-β balance; COL6A6^+^ fibroblasts reinforced ECM structure (COL6A6/COL6A3) and inhibited Wnt signaling (SFRP2); MAB21L1^+^ fibroblasts suppressed SMAD signaling (through NR2F1 upregulation) and blocked cell migration. Metastatic niches exhibited significant depletion of protective fibroblast subpopulations (F2RL2^+^: reduced from 1.9% to 0.1%; COL6A6^+^: reduced from 3.1% to 1.3%); this depletion coincided with ECM dissolution and disrupted spatial coordination of immune markers (proximity-dependent HLA-DRA/CXCL9 expression). High expression of these DEGs correlated with improved recurrence-free survival (e.g., COL6A6: HR = 0.19, p = 0.002); genomic deletions of the DEGs were significantly more frequent in metastases versus primary tumors (e.g., MAB21L1 deep deletion: 9.13% vs. 0.57%).

**Conclusion:** An unbiased transcriptomic approach revealed specialized fibroblast subpopulations as central gatekeepers against LNM via ECM, immune, and signaling barriers. Their erosion drives stromal collapse in metastasis. Epigenetic modulators (Valproic Acid, Vorinostat) were prioritized for therapeutic restoration of suppressive stroma.

## Introduction

Breast cancer metastasis to regional lymph nodes (LNs) is a critical prognostic indicator and driver of systemic dissemination [1,2]. It involves a complex multistage process, in which cells and other factors from primary tumors escape primary sites, travel via lymphatic circulation, prime premetastatic niches in LNs, and seed in LN tissues [3]. Components of the tumor microenvironment (TME)—including cancer-associated fibroblasts (CAFs), extracellular matrix (ECM), and immune cells—orchestrate metastatic progression [4,5]. While some CAFs facilitate metastasis, emerging evidence suggests specialized fibroblast subpopulations in primary tumors may establish metastasis-constraining stromal barriers [6,7], potentially through ECM fortification, immune coordination, and signaling modulation. However, comprehensive molecular drivers of stromal reprogramming in LN metastasis—particularly those involving spatial disorganization and ECM dissolution—remain poorly defined. The heterogeneity, plasticity, and interplay of stromal components in metastatic niches are unresolved, creating a need for multi-omics approaches including new technologies like scRNA-seq and spatial transcriptomics [8].

Here, we integrate bulk, single-cell, and spatial transcriptomics to systematically define conserved molecular alterations in breast cancer LN metastasis. We aimed to: (1) identify consistently dysregulated genes (DEGs) in LN metastases across independent cohorts; (2) resolve their cell-type specificity and spatial architecture; (3) characterize functional reprogramming in metastatic niches; and (4) evaluate their prognostic/therapeutic relevance. Critically, this approach was agnostic to cell-type mechanisms; fibroblast involvement was investigated *after* DEG identification.

## Methods

### Differential Expression Analysis

To identify genes with altered expression in Lymph node metastasis, we analyzed three independent Bulk RNA-seq datasets (GSE110590, GSE193103, GSE209998) comparing primary breast tumors to metastatic lesions in the lymph node metastases (LNMs). Raw count matrices were processed through DESeq2 (v1.38.3) in R (v4.2.2) to pinpoint differentially expressed genes (DEGs). The full code for this analysis can be obtained from the corresponding author upon request.

For the GSE193103 and GSE209998 datasets, we implemented a two-stage workflow:

1. **Tissue-Specific Gene Exclusion**: By comparing normal Lymph node and breast tissues (adjusted *p* < 0.05), we identified and removed genes with organ-specific expression to reduce confounding signals. In contrast, the GSE110590 dataset lacked matched normal tissue data, preventing tissue-specific filtering.
2. **Primary vs. Metastasis Comparison**: Lymph node metastasis samples were directly compared to primary breast tumors using a ∼Group design matrix, with primary tumors designated as the baseline

We applied likelihood ratio tests with independent filtering and stabilized log2 fold changes (LFC) using the ashr method to refine effect size estimates. DEGs were defined as genes with adjusted *p* < 0.05 and |LFC| >1 (a 2-fold change threshold).

To isolate consistent BCLM-associated genes, we overlapped DEGs from all three datasets using a Venn diagram approach, identifying 11 consensus genes. From these, three genes were prioritized for deeper investigation. Selection criteria included Consistency (Robust differential expression (significant *p* and LFC) across all datasets) and Novelty (Underexplored roles in Lymph node metastasis biology).

### Single-Cell RNA-Seq Data Processing and Analysis

Single-cell RNA sequencing (scRNA-seq) data of primary tumors and paired metastatic lymph nodes in four breast cancer patients from the public dataset GSE225600 (10X Genomics platform) was processed using Seurat (v5.0). Raw gene expression matrices were loaded with Read10X, and cell barcodes were matched to metadata from the original study to retain only high-confidence annotations using exact string matching with regular expression processing (e.g., converting “L2_AAAAAAAA” to “AAAAAAAA-L2”). Quality control excluded cells with <200 or >6,000 detected genes or >25% mitochondrial content, followed by normalization (LogNormalize; scale factor = 10,000) and identification of 3,000 highly variable genes. Dimensionality reduction was performed via PCA (top 50 PCs), and cells were clustered using a shared nearest-neighbor graph (FindNeighbors, k=20) and Louvain algorithm (FindClusters, resolution=0.5). Cell types were assigned based on pre-existing metadata to ensure consistency with the original study.

For cross-sample integration, Harmony (theta=2) was applied to correct for patient-specific batch effects using the RunHarmony function (grouping variable: sample_id, dimensions=1:20). Differential expression analysis between tissue compartments was performed with FindMarkers (Wilcoxon rank-sum test; min.pct=0.25, log2FC threshold=0.25) with Benjamini-Hochberg correction. Geneset enrichment analysis used clusterProfiler (v4.0) with GO Biological Processes ontology and human genome annotations (org.Hs.eg.db).

Fibroblast subpopulations were isolated for focused analysis using subset(x = sc, celltype == “Fibroblast”), followed by subclustering (resolution=0.8) and label transfer to the main object. DEG-positive cells were defined using normalized expression >0 in the RNA assay and characterized through Prevalence analysis across tissue types, Subcluster-specific expression profiling, Pathway enrichment in DEG+ vs DEG- subpopulations, and Tissue-specific differential expression within DEG+ fibroblasts.

Visualizations were generated with ggplot2 (v3.4.0), including: UMAP embeddings (DimPlot) colored by tissue type, cell type, and DEG status, Violin plots (VlnPlot) with boxplot overlays, Dot plots (DotPlot) of top markers, Co-expression networks (cnetplot) of enriched pathways, Enhanced volcano plots (EnhancedVolcano) for DEG results. All analyses used R (v4.3.0) with Bioconductor (v3.17) packages including Seurat, SingleCellExperiment, and org.Hs.eg.db.

### Spatial Transcriptomics Analysis of Metastasis-Associated Genes in Primary Breast Cancer

Spatial transcriptomics data from four primary breast tumors (T2, T3, T6, T7) from the public dataset GSE225600 were processed and analyzed to resolve the spatial architecture of metastasis-associated genes. Raw Visium data (10x Genomics) underwent quality control, retaining only tissue-embedded spots using Seurat’s ‘subset’ function based on ‘in_tissuè metadata. We converted raw matrices to HDF5 format via ‘DropletUtils::write10xCounts’ and loaded spatial data using ‘Seurat::Load10X_Spatial’, ensuring barcode alignment with tissue positions through manual metadata integration. Sequencing depth was visualized using ‘SpatialFeaturePlot’ for nCount_Spatial, confirming absence of low-quality spots (<5K UMIs) and revealing patient-specific distributions.

Cell type deconvolution employed RCTD (spacexr) with patient-matched scRNA-seq references. Single-cell data were subset to primary samples (T2, T3, T6) using sample-specific suffixes, normalized via ‘SCTransform’, and converted to ‘SummarizedExperiment’ objects. Spatial data were converted to ‘SpatialExperiment’ format, and RCTD was executed in multi-mode (max_cores=20, max_multi_types=3) to generate cell type weight matrices. Cell type distributions were visualized spatially by integrating weights with spatial coordinates and plotting with ‘ggplot2’.

Spatial autocorrelation of metastasis-associated genes (MAB21L1, F2RL2, COL6A6) was quantified using Moran’s I. A k-nearest neighbor graph (k=5) was constructed from spatial coordinates via ‘spdep::knn2nb’, and significance was tested with ‘moran.test’ (α=0.05). Genes were considered clustered if Moran’s I >0 with adjusted p<0.05.

Microenvironmental associations were assessed through spatial proximity analysis. For Immune/tumor niches definition, gene expression (CD3D, EPCAM, CD68, KRT19, LYZ, STAB1, PDCD1, CTLA4, CD8A, CD163, MARCO) was extracted from spatial data assays, and cell type weights (epithelial, myeloid, T cell) were converted from RCTD results. Three niches were defined using quantile thresholds on non-zero expression: The Tumor niche required (EPCAM > 60th percentile or KRT19 > 55th percentile or epithelial weight >0.5) while excluding high myeloid/T-cell markers; The Myeloid niche required (CD68 > 55th percentile or LYZ > 55th percentile or [STAB1 > 75th percentile & myeloid weight >0.4]) while excluding epithelial/T-cell markers; The T-cell niche required (CD3D > 55th percentile or CD8A > 55th percentile or [PDCD1 > 65th percentile & CTLA4 >0] or T-cell weight >0.4) while excluding epithelial/myeloid markers. Distances between gene-positive spots (expression >0) and niche centroids were calculated using ‘FNN::get.knnx’ (k=1). Permutation tests (n=1000) compared observed median distances to random spot distributions, with Benjamini-Hochberg correction. Proximity enrichment was calculated as the percentage of gene-positive spots within 100µm of niches versus random expectation. Marker correlations used Spearman’s ρ on ‘FetchDatà-extracted expression.

Functional crosstalk was characterized by differential expression analysis of gene-positive spots within 100µm versus >500µm from myeloid hotspots (top 10% LYZ+ spots). DEGs were identified via ‘FindMarkers’ (logFC>0.25, p<0.05) and functionally annotated using ‘clusterProfiler::enrichGÒ (Biological Process, org.Hs.eg.db). Pathway significance required adjusted p<0.05. Spatial expression of immune-activation genes (e.g., HLA-DRA, CXCL9) was visualized in fibroblast subsets using ‘SpatialFeaturePlot’. All analyses used R (v4.3.2) with Seurat (v5.1.0), spacexr (v1.3.0), and spatial dependencies (spdep, spatstat). Analyses used R (v4.1.0), Seurat (v4.0), spacexr, spdep, and clusterProfiler, with visualization via ggplot2, SpatialFeaturePlot, and pheatmap; adjusted p-values <0.05 were considered significant throughout.

#### Clinical Validation

##### Genetic Alteration Profiling

We interrogated genetic alterations in the DEGs using cBioPortal (https://www.cbioportal.org/). Metastatic cohorts included Metastatic Breast Cancer (DFCI, Cancer Discov 2020), Metastatic Breast Cancer (INSERM, PLoS Med 2016), The Metastatic Breast Cancer Project (Archived, 2020), and The Metastatic Breast Cancer Project (Provisional, December 2021), while primary cohorts derived from following datasets Breast Cancer (METABRIC, Nature 2012 & Nat Commun 2016), Breast Invasive Carcinoma (British Columbia, Nature 2012), Breast Invasive Carcinoma (Broad, Nature 2012), Breast Invasive Carcinoma (Broad, Nature 2012), and Breast Invasive Carcinoma (TCGA, Firehose Legacy). Copy-number alterations (CNAs) were detected via GISTIC 2.0 and DNAcopy, with amplification/deletion frequencies calculated as the percentage of samples harboring alterations per gene. RNA-seq expression trends (e.g., **F2RL2** downregulation in metastases) were cross-referenced with dominant CNAs (amplifications for upregulated genes, deletions for downregulated) to infer drivers of metastatic adaptation.

##### Progression-Associated Expression Dynamics

The expression patterns of DEGs were evaluated across breast cancer progression stages using the TNMplot platform (https://tnmplot.com/analysis/). RNA-seq data from the Breast Cancer RNA-seq cohort—comprising 113 normal breast tissues, 1,097 primary tumors, and 7 metastatic lesions—were analyzed. For each gene, median expression values were calculated, and fold changes (Fc) were determined for tumor versus normal (T/N) and metastatic versus tumor (M/T) comparisons. A Kruskal-Wallis test (P < 0.05) assessed overall expression differences among groups, followed by Dunn’s post-hoc test with Benjamini-Hochberg correction for pairwise comparisons (normal vs. tumor, tumor vs. metastatic). Statistical significance, fold changes, and expression distributions were visualized using TNMplot-generated box plots. Non-parametric methods were selected due to non-normal data distribution and the small metastatic sample size, ensuring robust detection of genes suppressed or sustained during metastasis.

##### Kaplan-Meier Survival Analysis

Using the KM Plotter tool (https://kmplot.com/analysis), we evaluated the prognostic relevance of the six DEGs in 2,976 primary breast tumors. Patients were stratified into “high” and “low” expression groups via the tool’s Auto Select Best Cutoff feature, which algorithmically identifies thresholds that maximize survival differences. The analysis included data from up to 10 years (120 months) of follow-up. Statistical significance was assessed with log-rank tests (p<0.05 threshold), and Cox proportional hazards models calculated hazard ratios with corresponding 95% confidence intervals. This linked BC LNMs-associated expression patterns (e.g., F2RL2 downregulation) to survival outcomes in early-stage disease.

##### Computational Identification of Therapeutic Candidates via the Comparative Toxicogenomics Database

Candidate therapeutic agents were systematically identified using the Comparative Toxicogenomics Database (CTD) through the Drug Signatures Database (DSigDB), with a focus on human-specific gene interactions. Selection criteria included any increases or effects on chemical gene interactions. For more targeted candidate selection, environmental pollutants (e.g., Benzo(a)pyrene, Particulate Matter) were excluded, and non-FDA-approved compounds were retained only if backed by clinical trial evidence in oncology (e.g., Resveratrol). Prioritization criteria were two in total: (1) number of CTD supportive references, and (2) relevance to metastatic mechanisms.Shortlisted molecules were also evaluated for biological relevance to breast cancer, marking their potential roles in proliferation inhibition, immune modulation, or epigenetic modification. This analysis is a computational screen; experimental validation was not within the purview of this work, but findings provide a prioritized hypothesis for metastasis-targeted therapy functional study in the future.

## Results

### 1. Conserved Fibroblast-Specific Gene Downregulation in Lymph Node Metastasis Integration of 3 RNA-seq cohorts identifies consistent downregulation of MAB21L1, F2RL2, and COL6A6 in LNMs (Table 1)

#### 1.1 Bulk transcriptomics identifies metastasis-suppressing genes

To define transcriptomic drivers of breast cancer lymph node metastasis (LNM), we integrated three bulk RNA-seq datasets. Transcriptomic analysis of three independent datasets (GSE110590, GSE193103, and GSE209998) revealed consistent downregulation of MAB21L1, F2RL2, and COL6A6 in LNMs compared to primary breast tumors (Table 1). All three genes met stringent statistical thresholds (unadjusted p < 0.01, adjusted p < 0.05; DESeq2 LRT) with uniform directionality across datasets. MAB21L1 exhibited the most pronounced down regulation in the GSE110590 dataset (log2FC = −3.19), while COL6A6 showed the strongest suppression in GSE193103 (log2FC = −2.80). F2RL2 demonstrated robust down regulation across all datasets, with the largest effect in GSE209998 (log2FC = −3.69). (Table 1) These bulk findings prompted cell-type-resolved investigation.

**Table 1:**
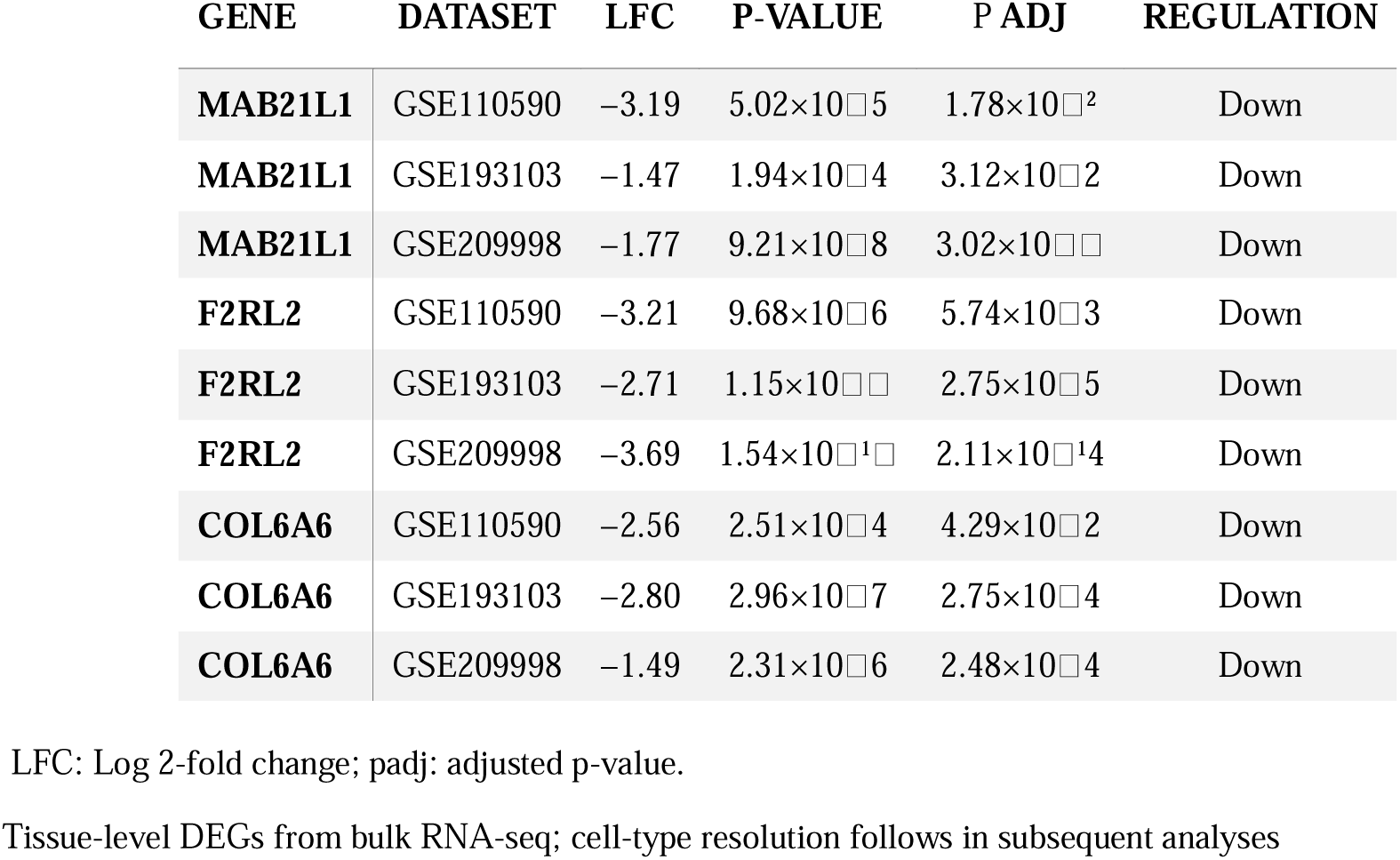
Consistent Dysregulation of three DEGs in lymph node metastasis from breast cancer.

#### 1.2 Fibroblasts are the exclusive cellular reservoir depleted in metastases (scRNA-seq)

To resolve cellular specificity, we analyzed single-cell transcriptomes of primary tumors and matched LNMs. Single-cell RNA sequencing analysis of primary tumors and matched lymph node metastases (GSE225600, n=4 patients) revealed distinct transcriptional landscapes between tissues. Figure 1A shows UMAP visualization of transcriptional differences between primary tumors and LNMs, revealing distinct tissue-specific clustering. Figure 1B demonstrates these patterns persist across patients, confirming biological consistency. Figure 1C displays a heatmap of 15+ canonical cell-type markers (EPCAM, PECAM1, CD3D, etc.) with z-score normalized expression (blue-low to red-high) to validate annotation quality across all cells. Quantitative analysis in Figure 1D confirmed compositional differences: LNMs were lymphocyte-rich (T cells: 57.6%; B cells: 23.1%) while primary tumors showed stromal enrichment (endothelial cells: 38.1%; epithelial cells: 15.6%).

**Figure 1.**
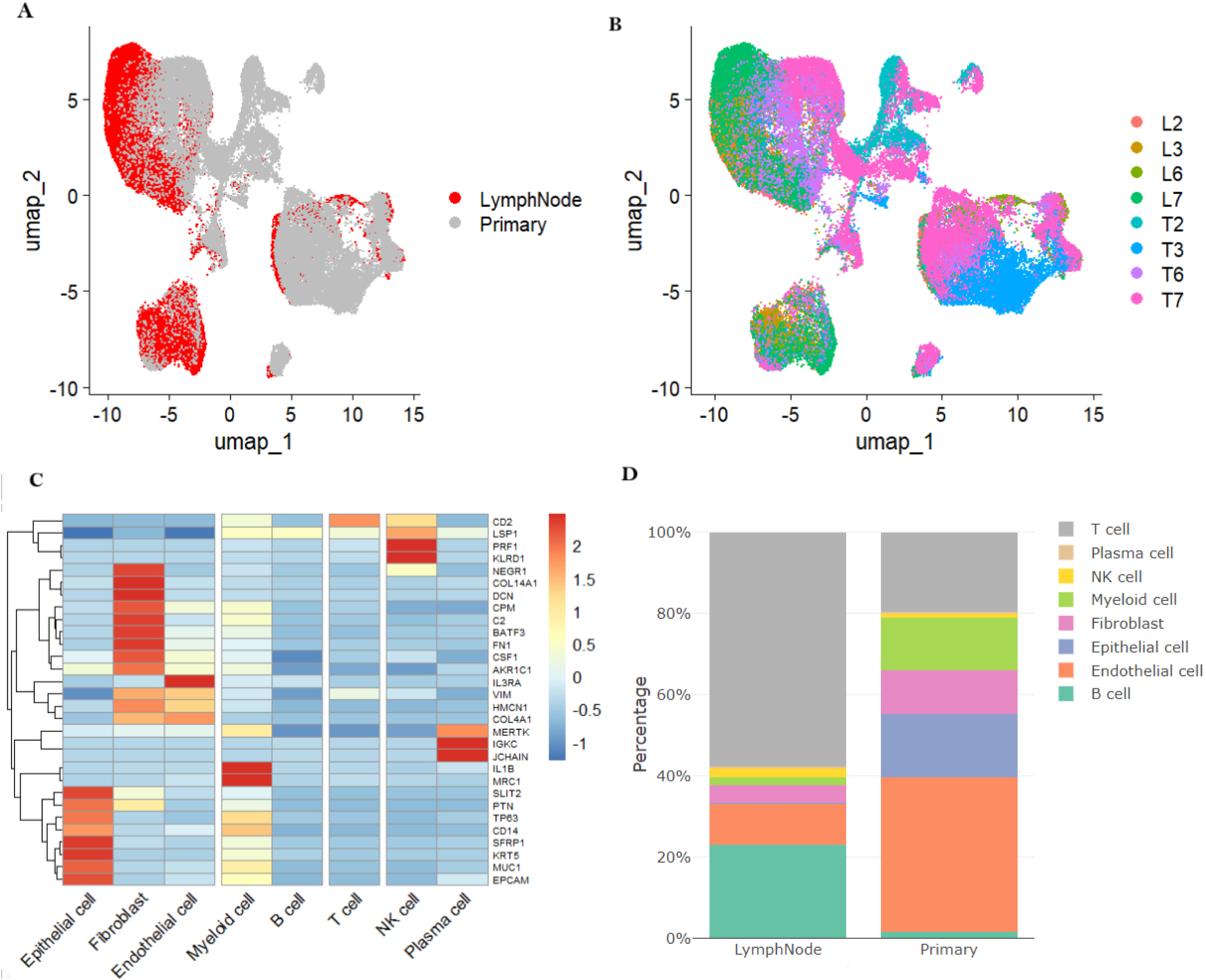
Single-cell landscape of primary breast tumors and matched lymph node metastases (LNMs). (A) UMAP projection colored by tissue origin (Primary tumor vs. LNM). (B) UMAP colored by patient ID, showing inter-individual variation. (C) Heatmap of annotation markers only (rows = genes, columns = cells) used for cell type validation, Color scale: blue (low) to red (high) z-score normalized expression. (D) Stacked bar plot of cell type frequencies. Data source: GSE225600.

Spatial patterns of specific markers in LNMs were assessed to confirm their metastatic (not normal lymph node) identity. Supplementary Figure 1 establishes the metastatic identity of LN tissues through pan-tissue VIM expression, confirming near-universal mesenchymal/stromal activation (present in most of cells). While epithelial markers (EPCAM/CDH1/KRT19) and MKI67 show spatially restricted, low-frequency expression (consistent with metastatic dissemination), their non-random spatial segregation from VIM-high zones (arrows) excludes normal LN contamination. Critically, the co-localization of residual epithelial markers with proliferative (MKI67+) niches supports true metastatic deposits amid stromal dominance. These spatial features—stromal dominance, constrained epithelial marker expression, and proliferative epithelial-stromal co-localization—collectively validate the profiled tissues as metastatic lesions exhibiting expected EMT dynamics, ensuring fidelity for subsequent metastatic niche analyses.

To resolve the cellular origin of conserved DEGs identified in bulk analyses, we assessed cell-type-specific expression patterns of MAB21L1, F2RL2, and COL6A6. Expression of the metastasis-associated genes MAB21L1, F2RL2, and COL6A6 is cell-type-specific in both primary tumors and lymph node metastases (LNMs), as demonstrated by distinct expression patterns across diverse cell populations. Notably, fibroblasts exhibited robust enrichment for all three genes compared to other major cell types, showing consistently higher expression levels and a greater fraction of expressing cells in both tissue contexts. (Figure 2A-C) Fibroblasts in primary tumors exhibited significantly higher proportions of DEG-positive cells compared to fibroblasts in LNMs (e.g., F2RL2: 1.9% vs. 0.1%; Figure 2D & Supplementary Table 1 & Supplementary Figure 2). Minimal expression was observed in non-fibroblast lineages. This fibroblast-specific expression pattern and metastatic depletion establish fibroblasts as the primary cellular reservoir for these metastasis-suppressing factors, directing focus to fibroblast subpopulation dynamics for subsequent mechanistic analyses.

**Figure 2.**
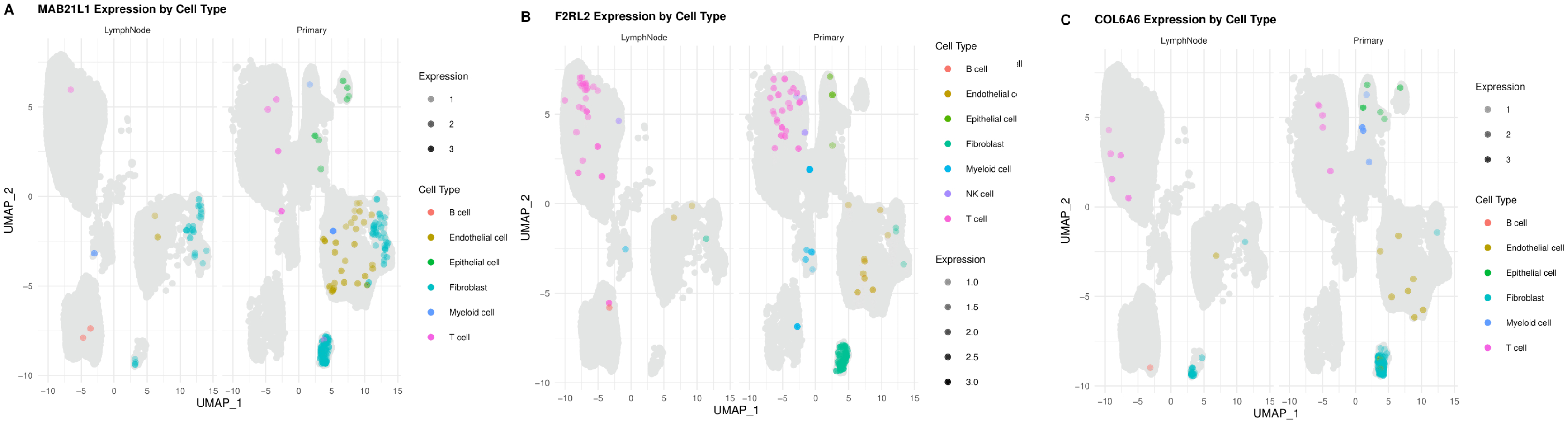
Expression of the metastasis-associated genes (MAB21L1, F2RL2, COL6A6) is cell-type-specific in both primary tumors and LNMs. Fibroblasts in primary tumors show elevated expression of these genes compared to fibroblasts in LNMs

### 2. Functional Specialization of Fibroblast Subpopulations in Primary Tumors

#### 2.1 Fibroblast subclusters orchestrate complementary protective roles

To determine whether DEG-expressing fibroblasts represent functionally distinct subpopulations, we performed sub clustering and pathway enrichment analysis. In primary breast cancer, DEGs were predominantly localized to specific fibroblast subpopulations (clusters 1, 7, 8), where COL6A6 showed maximal prevalence in cluster 7 (19.1% of cells). (Figure 3A&B, Supplementary Table 2) Enrichment analysis revealed distinct functional specialization among fibroblast subpopulations expressing the DEGs(*Figure 3C)*: Subcluster 1, characterized by concurrent high expression of *F2RL2* (8.59), *MAB21L1* (8.30), and *COL6A6* (10.07), (Figure 3A) was enriched for protein homeostasis pathways (chaperone-mediated folding, PERK-mediated unfolded protein response) and immune/stress regulation (chronic inflammatory response, NOD2 signaling), indicating a role in maintaining proteostatic resilience. Subcluster 7, exhibiting the highest *COL6A6* expression, showed dominant enrichment for extracellular matrix (ECM) organization (collagen fibril assembly, basement membrane formation), TGF-β/BMP signaling, wound healing, and immune modulation (complement activation), highlighting its multifunctional role in stromal remodeling and immune coordination. Subcluster 8, with prominent *COL6A6* but moderate overall expression, was linked to mitochondrial energy metabolism (oxidative phosphorylation, ATP synthesis) and inflammatory microenvironment modulation (respiratory burst, leukocyte migration), suggesting metabolic support functions. (Figure 3C) Critically, these subclusters represent non-overlapping, specialized fibroblast states supporting complementary pro-tumor processes. Their collective downregulation in LNMs creates a permissive metastatic niche by simultaneously disrupting: (i) physical ECM barriers (Subcluster 7), (ii) stress-defense systems (Subcluster 1), and (iii) metabolic/immune coordination (Subclusters 7 & 8). This fibroblast-driven stromal collapse aligns with a mechanistic model where loss of compartmentalized stromal functions enables immune evasion and metastatic progression (Figure 3C, Supplementary Tables SC 3 & 4).

**Figure 3.**
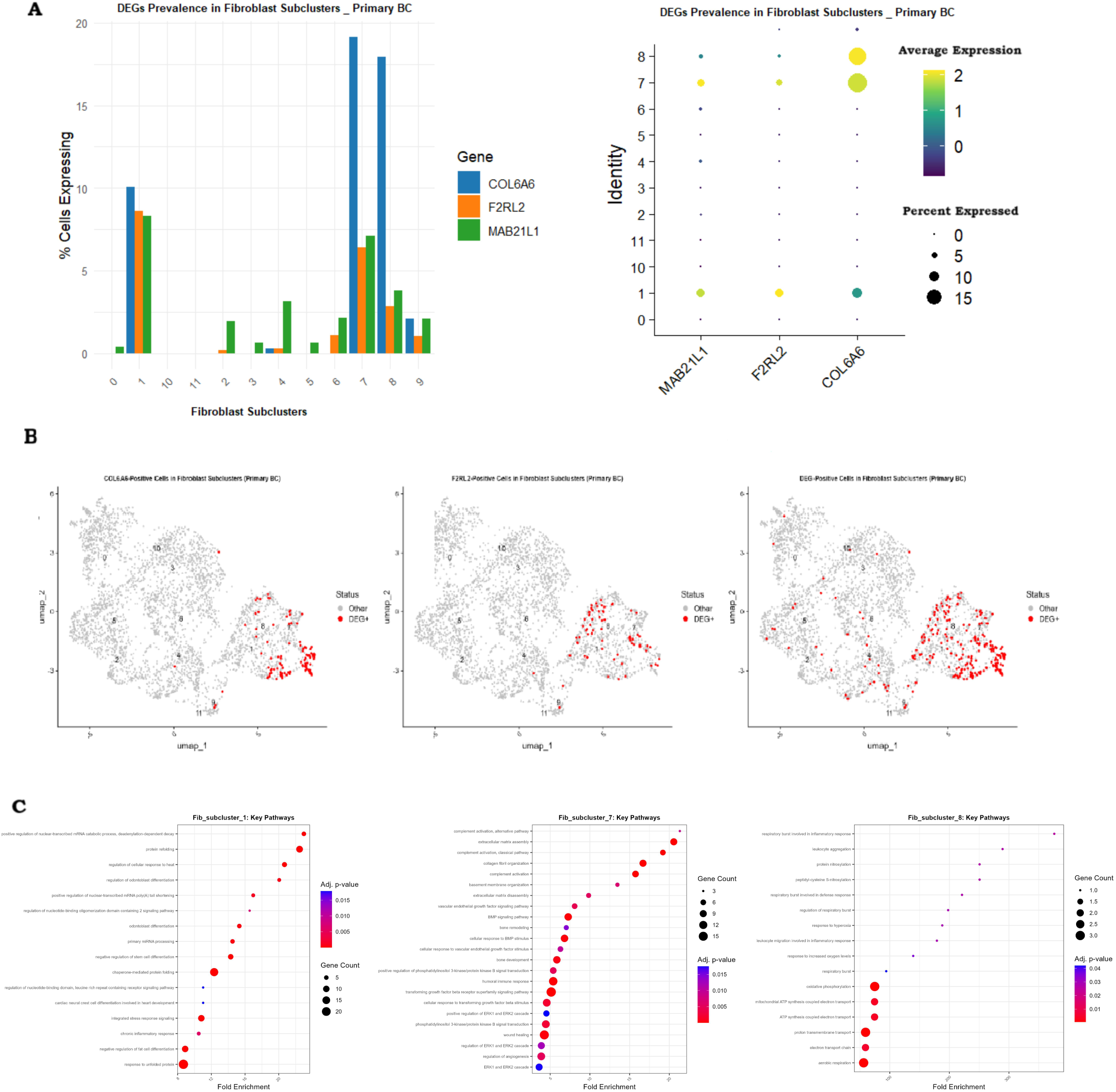
(A&B) In primary breast tumors, DEGs were localized to specific fibroblast subclusters, with COL6A6 highly expressed in cluster 7. (C) Functional analysis revealed distinct specialization - subcluster 1 in proteostasis, subcluster 7 in ECM remodeling/immune coordination, and subcluster 8 in metabolism/inflammation (hypergeometric testing and BH correction. via clusterProfiler)

#### 2.2 Pathway-defined defense mechanisms against metastasis

To define the functional mechanisms by which fibroblast subpopulations constrain metastasis, we performed pathway enrichment and marker validation for each DEG-specific subset. F2RL2+ fibroblasts coordinate a sophisticated defense network against metastasis through three synergistic mechanisms, validated by robust marker-pathway integration. (Figure 4 A-D) First, extracellular matrix reorganization is driven by structural regulators THBS4 (log2FC=4.85, padj=2.46e-31), CILP (log2FC=3.29, padj=9.64e-33), and COL6A3 (log2FC=2.06, padj=4.66e-21), which orchestrate collagen fibril assembly while MMP11 (log2FC=3.09, padj=8.71e-07) enables controlled matrix disassembly. (Figure 4 B) Second, TGF-β/BMP signaling modulation is evidenced by concurrent activation (INHBA↑, log2FC=1.88, padj=5.94e-07) and suppression (FSTL1↑, log2FC=1.42, padj=6.00e-12), creating a balanced signaling environment that limits EMT progression. (Figure 4 A&B) Crucially, third, immune-metastasis crosstalk is mediated through CXCL14 (log2FC=3.26, padj=1.23e-17)-driven monocyte chemotaxis and C1S (log2FC=1.49, padj=2.34e-09)-initiated complement activation, recruiting tumor-suppressive leukocytes while TNFSF10 (log2FC=1.67, padj=3.62e-10) primes T-cell responses. (Figure 4 B) These processes are unified by potent metastasis suppression via SFRP2 (log2FC=2.18, padj=3.17e-31)-mediated Wnt inhibition and FSTL1-dependent BMP constraint, collectively establishing F2RL2+ fibroblasts as conductors of a multi-layered defense that integrates ECM fortification, immune recruitment, and pathway blockade to contain dissemination.

**Figure 4.**
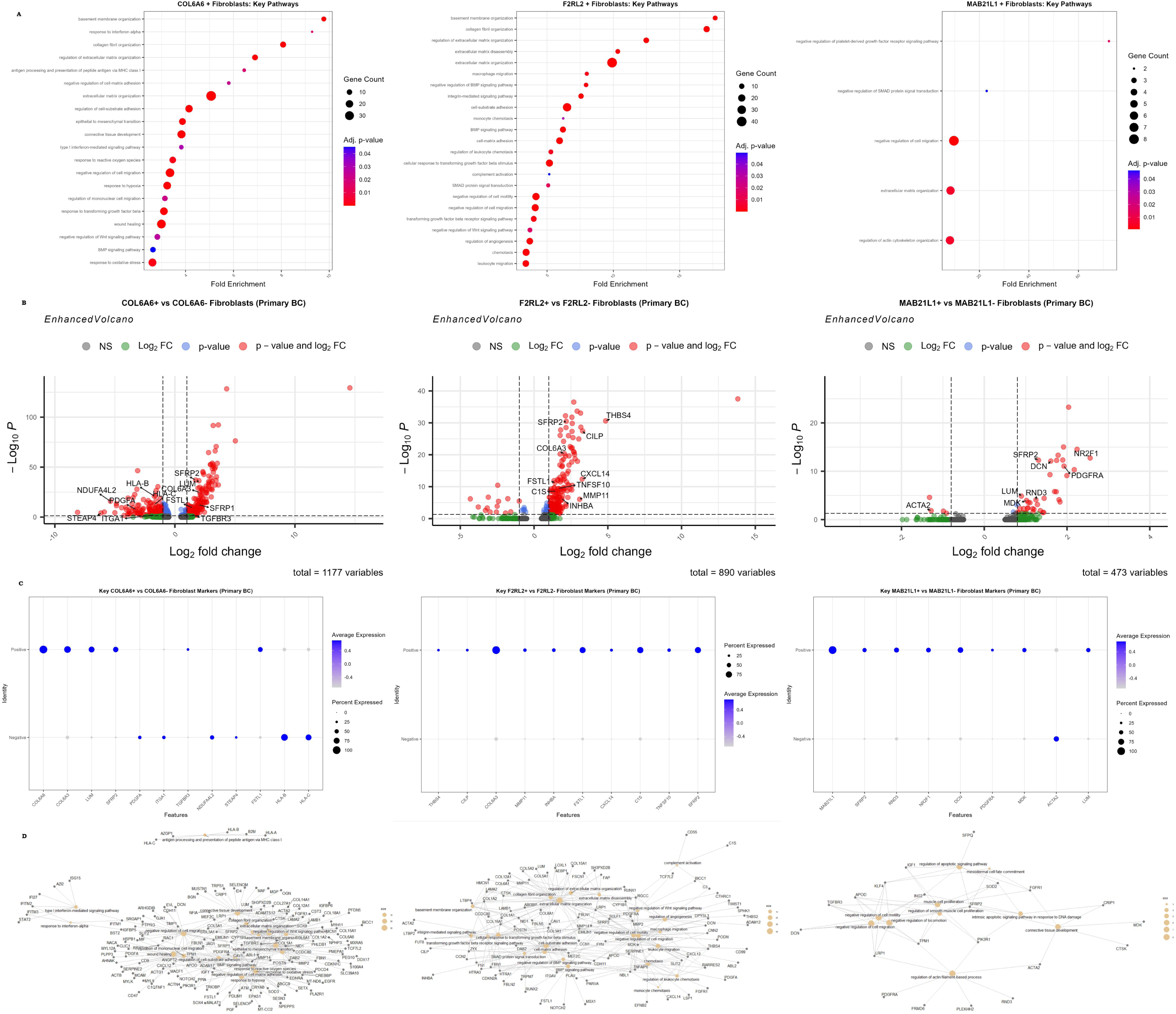
Molecular pathways and expression signatures in fibroblast subpopulations of primary breast cancer. Analysis of three fibroblast subtypes (COL6A6+, F2RL2+, MABEL1+) reveals key pathways associated with metastasis suppression. Differential gene expression (log fold change) and statistical significance (adjusted p-value) are shown for critical pathways. Enhanced Volcano plots highlight genes involved in extracellular matrix (ECM) regulation, TGF-β/BMP signaling, and immune mediation. Fold change analyses validate functional enrichment in ECM remodeling, signaling inhibition, and immune recruitment, supporting an anti-metastatic defense network. Gene counts and adjusted p-values quantify pathway significance. (hypergeometric testing and BH correction. via clusterProfiler)

COL6A6+ fibroblasts deploy a multi-layered defense against metastasis through seven synergistic pathways, each validated by robust molecular markers. (Figure 4 A-D) Central to this defense is extracellular matrix fortification, driven by dominant upregulation of structural genes COL6A6 (log2FC=14.58), COL6A3 (log2FC=2.07), and LUM (log2FC=2.01), which collaboratively establish a physical barrier against invasion. This ECM reinforcement is coupled with potent migration suppression via SFRP2 (log2FC=1.72), a Wnt inhibitor that blocks pro-dissemination signals, and downregulation of motility drivers PDGFA (log2FC=-3.27) and ITGA1 (log2FC=-1.72). (Figure 4 A&B) Crucially, EMT constraint is mediated through TGFBR3 (log2FC=1.41), which sequesters TGF-β to limit epithelial-mesenchymal transition, while hypoxic adaptation is evidenced by suppression of HIF-1α targets NDUFA4L2 (log2FC=-5.37) and STEAP4 (log2FC=-6.16). Further metastasis containment arises from Wnt pathway inhibition via SFRP1 (log2FC=2.00) and TGF-β/BMP modulation through FSTL1 (log2FC=1.47), with immune crosstalk facilitated by HLA-B/C downregulation (log2FC≈-1.70) attenuating antigen presentation. (Figure 4 A&B) Collectively, these marker-validated pathways position COL6A6+ fibroblasts as architects of a multi-layered stromal defense system that integrates structural fortification, signaling interception, and metabolic reprogramming to constrain metastatic progression.

Functional enrichment analysis of MAB21L1+ fibroblasts in primary breast tumors revealed key pathways implicated in lymph node metastasis suppression. (Figure 4 A-D) Critically, we identified negative regulation of cell migration (padj < 0.05), validated by significant upregulation of motility inhibitors SFRP2 (log2FC=1.31, padj=5.27e-13) and DCN (log2FC=1.61, padj=7.76e-13). Equally compelling was the enrichment for negative regulation of SMAD protein signal transduction, mechanistically supported by overexpression of NR2F1 (log2FC=2.24, padj=2.88e-15), a known TGF-β pathway suppressor that inhibits epithelial-mesenchymal transition. The negative regulation of platelet-derived growth factor receptor signaling pathway was corroborated by concomitant upregulation of PDGFRA (log2FC=1.93, padj=8.76e-12) – which may act as a decoy receptor – paired with downregulation of its activator MDK (log2FC=1.02, padj=0.0001). (Figure 4 A&B) Furthermore, regulation of actin cytoskeleton organization was evidenced by elevated expression of cytoskeletal remodelers RND3 (log2FC=1.25, padj=8.12e-05) and reduced ACTA2 (log2FC=-1.28, padj=0.014), indicating *restricted* cellular motility. Collectively, these pathway-markers align functionally to constrain metastasis through: 1) ECM stabilization (DCN, LUM), 2) blockade of pro-metastatic signaling (NR2F1, SFRP2), and 3) inhibition of stromal-tumor crosstalk (PDGFRA). The specialization of these mechanisms reveals how coordinated fibroblast functions create layered protection against dissemination, whose erosion in metastasis enables multi-faceted stromal collapse.

#### 2.3 Stromal Collapse in Metastatic Niches

To characterize how metastatic progression alters protective fibroblast functions, we compared DEG-expressing subpopulations in primary tumors versus lymph node metastases (LNMs). Single-cell profiling revealed three coordinated fibroblast alterations driving LN metastasis: (1) selective depletion of tumor-suppressive F2RL2+ (0.1% vs 1.9% primary) and COL6A6+ (1.3% vs 3.1%) subsets. Residual cells (i.e., the rare F2RL2+ or COL6A6+ fibroblasts persisting in LNMs despite significant depletion of these subsets in metastatic niches) showing pathogenic reprogramming including upregulation of lymphoid chemokines (“CCL19/21” in F2RL2; log FC ≈ +4.75) and ion transporters (*SLC26A7* in COL6A6; log FC = +7.50). (2) ECM dissolution via loss of structural programs (*COL3A1*, *DCN* in F2RL2; *COL6A6*, *FBLN5* in COL6A6; log FC ≤ −2.62); and (3) In persistent MAB21L1 fibroblasts (1.8% vs 2.7% in LNMs vs Primary BC), non-significant trends suggested metabolic rewiring (↓DES/ACTG2; log FC ≈ −6.3) and adhesion remodeling (↑*MUCL1*; log FC = +5.48). This stromal collapse—marked by ECM degradation, immune dysregulation, and metabolic shifts—creates a permissive niche for metastasis. (Supplementary Table 5)

### 3. Spatial Architecture of Fibroblast Defense Niches

To resolve the spatial organization of metastasis-suppressing fibroblast niches, we first established data quality and baseline microenvironment architecture. Spatial QC revealed T3 had the highest sequencing depth (∼60K UMIs, epithelial hotspots), while T7 showed the lowest (∼30K UMIs, stromal zones). All samples exhibited right-skewed nCount distributions, with most spots capturing moderate RNA (10K–30K UMIs; mixed cell types) and rare high-count outliers (>40K; epithelial/myeloid foci). (Figure 5A) This robust data quality supported three distinct patterns: T2 displayed a unique fibroblast-myeloid reciprocal architecture (mutually exclusive high-weight zones) with sparse B/T cell presence. In contrast, T3/T6/T7 showed epithelial dominance, where fibroblast/myeloid weights increased only in epithelial-low regions. Strikingly, NK cells were universally excluded (weights ≈0), while T/B cells were nearly absent in T3/T6/T7. (Figure 5B) Critically, the absence of low-quality spots (<5K UMIs) confirms these lymphoid exclusion patterns reflect true biology rather than technical artifacts. (Figure 5A) The epithelial-fibroblast reciprocity in T3/T6/T7 suggests conserved tumor-stromal boundaries, while T2’s pattern implies distinct immune evasion mechanisms. This spatial heterogeneity provides essential context for interpreting DEG distribution patterns across individual metastatic defense niches.

**Figure 5.**
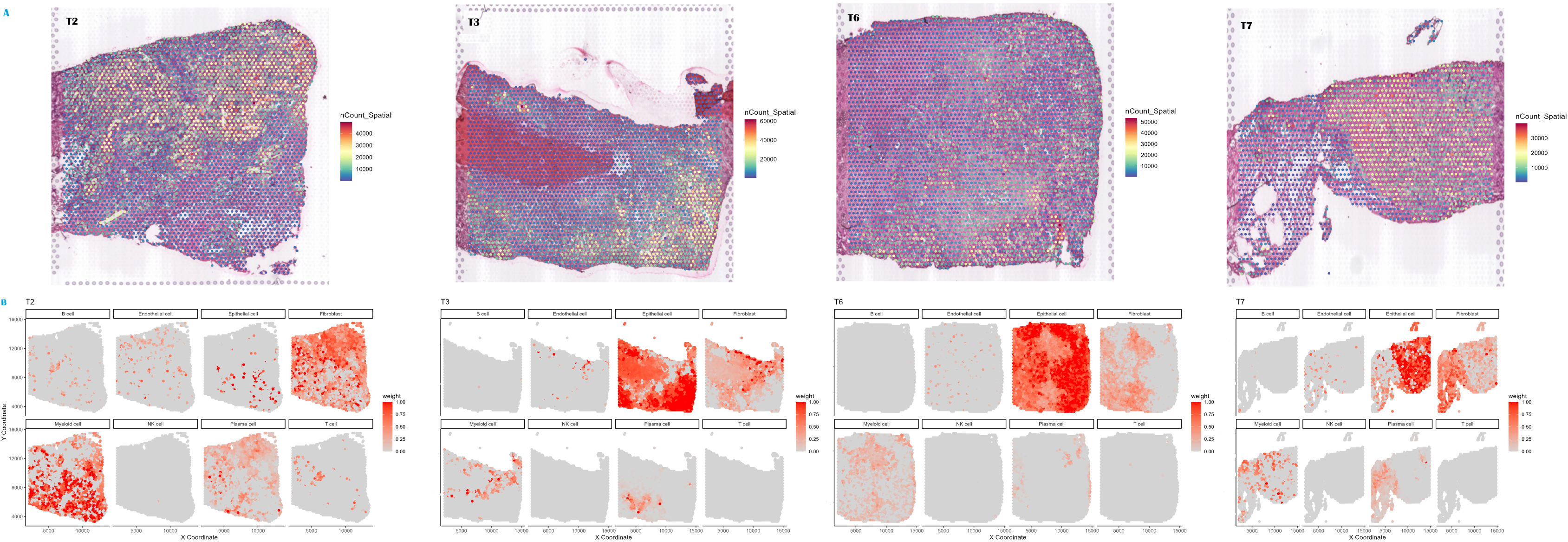
Spatial QC and tissue architecture. (A) nCount distributions: T3 highest depth (∼60K UMIs, epithelial foci), T7 lowest (∼30K, stromal). Most spots: 10K–30K UMIs (mixed cell types); outliers >40K (epithelial/myeloid). (B) Architectural patterns: T2 shows reciprocal fibroblast-myeloid exclusion zones. T3/T6/T7 exhibit epithelial dominance with fibroblast/myeloid restricted to epithelial-low regions. Universal NK exclusion; T/B cells absent in T3/T6/T7. Biological validity confirmed (no low-quality spots). Epithelial-fibroblast reciprocity (T3/T6/T7) indicates conserved tumor-stromal boundaries; T2 suggests distinct immune evasion. (descriptive QC and pattern visualization)

#### 3.1 Organized spatial patterning supports niche functions

To determine whether metastasis-suppressing fibroblast genes exhibit organized spatial distributions that could support coordinated defense functions, we quantified spatial autocorrelation patterns of MAB21L1, F2RL2, and COL6A6 across primary tumors. We characterized the spatial architecture of three metastasis-associated genes (*MAB21L1*, *F2RL2*, *COL6A6*) in four primary breast tumors using spatial transcriptomics, analyzing 3,153 spots in T2, 2,681 in T3, 3,510 in T6, and 1,793 in T7. (Supplementary Figure 3) Spatial autocorrelation revealed distinct patient-specific organization patterns. *F2RL2* exhibited significant clustering in T6 (Moran’s *I* = 0.018, p = 0.039) and T7 (*I* = 0.055, p = 0.0001) but random distribution in T2 (*I* = 0.008, p = 0.226) and T3 (*I* = 0.006, p = 0.286). *COL6A6* showed organized expression only in T2 (*I* = 0.020, p = 0.030), was undetectable in T3 and T6 due to extreme rarity (< 5 spots), and displayed random distribution in T7 (*I* = −0.012, p = 0.789). *MAB21L1* demonstrated clustering solely in T2 (*I* = 0.028, p = 0.005) with non-significant patterns elsewhere (T3 *I* = −0.0003, p = 0.160; T7 *I* = - 0.0007, p = 0.813) and undetectable expression in T6. This heterogeneity highlights variable spatial organization of molecular defenses across individuals. (Supplementary Table 6) These heterogeneous spatial patterns directly reflect the functional roles identified in single-cell analyses: organized F2RL2 hubs in T6/T7 enable coordinated ECM-immune crosstalk, structured COL6A6 in T2 forms physical barriers, and diffuse MAB21L1 supports metabolic functions. Critically, the near absence of structured COL6A6 organization in most tumors (3/4) reveals a common vulnerability in spatial defense systems that may license metastatic dissemination.

To determine how fibroblast subpopulation positioning enables metastasis-suppressive functions, we quantified spatial relationships between DEG-expressing fibroblasts and tumor microenvironment niches. F2RL2+ fibroblasts exhibited conserved immune-niche specialization in tumors T2, T6, and T7, showing significant myeloid proximity (BH-adj. P <0.001), and hotspot enrichment (12.9–15.4% vs. 9.6–10.2% random), along with correlations to myeloid (LYZ: ρ=0.044–0.099) and T-cell markers (CD3D: ρ=0.041–0.089). In contrast, F2RL2+ fibroblasts in T3 showed no significant spatial proximity to immune niches or marker correlations, suggesting context-dependent regulation(Supplementary Table 7 to 9, Supplementary Figure 4 A-D)COL6A6+ fibroblasts exhibited microenvironmental plasticity: tumor-adjacent in T3 (median distance=0µm; EPCAM: ρ=0.058), myeloid-proximal in T2/T6 (hotspot enrichment: 10.6–20.0%; T-cell proximity in T2: p =0), but epithelial-avoidant in T7 (KRT19: ρ=−0.081; no hotspot enrichment). Their scarcity in T3, T6, and T7 (COL6A6: T3 n=11, T6 n=5, T7 n=21) suggests erosion of tumor-restraining programs(Supplementary Table 7 to 9, Supplementary Figure 5A-D) MAB21L1+ fibroblasts demonstrated robust immune anchoring in **T3 (n=13), T6 (n=9), and T7 (n=10)**, with 11.1–20.0% of spots overlapping myeloid hotspots (p<0.001) and immune marker correlations (CD68: ρ=0.047 in T3; CD3D: ρ=0.040–0.051 in T3/T6). However, in **T2 (n=56)**, they lacked myeloid proximity and showed a negative correlation with LYZ (ρ=−0.056), indicating tissue-specific functional divergence driven by abundance shifts. (Supplementary Table 7 to 9, Supplementary Figure 6A-D)

These spatial patterns directly license specialized functions: precise positioning enables F2RL2 immune coordination, COL6A6 barrier formation, and MAB21L1 immune anchoring. Patient-specific disruption of these spatial relationships—through either mispositioning (T3 F2RL2) or depletion (COL6A6)—compromises metastasis containment by disconnecting fibroblast programs from their effector microenvironments.

#### 3.2 Proximity-dependent immune licensing

To test whether spatial proximity licenses functional immune activation by metastasis-suppressing fibroblasts, we analyzed gene expression changes in fibroblast subpopulations near (<100μm) versus far (>500μm) from myeloid niches. Critically, immune-activating programs were triggered exclusively when fibroblasts resided within <100 spatial units of myeloid cells, revealing a strict distance-dependent crosstalk mechanism.

F2RL2+ fibroblasts demonstrated conserved proximity-driven activation in T2 and T7 but functional erosion in T3/T6. In T2, F2RL2+ spots near myeloid cells showed robust upregulation of MHC-II machinery (HLA-DRA, P _adj_= 4.3e-14; CD74, P _adj_ = 4.2e-11) and interferon-response genes (CXCL9, P _adj_ = 7.8e-6; STAT1, P _adj_ = 2.7e-3), with pathway enrichment confirming antigen presentation via MHC class II (GO:0019886, P _adj_ = 8.0e-12), T-cell regulation (GO:0050870, P _adj_ = 8.5e-11), and leukocyte adhesion (GO:0007159, P _adj_ = 2.4e-8). (Figure 6 A-C)

**Figure 6.**
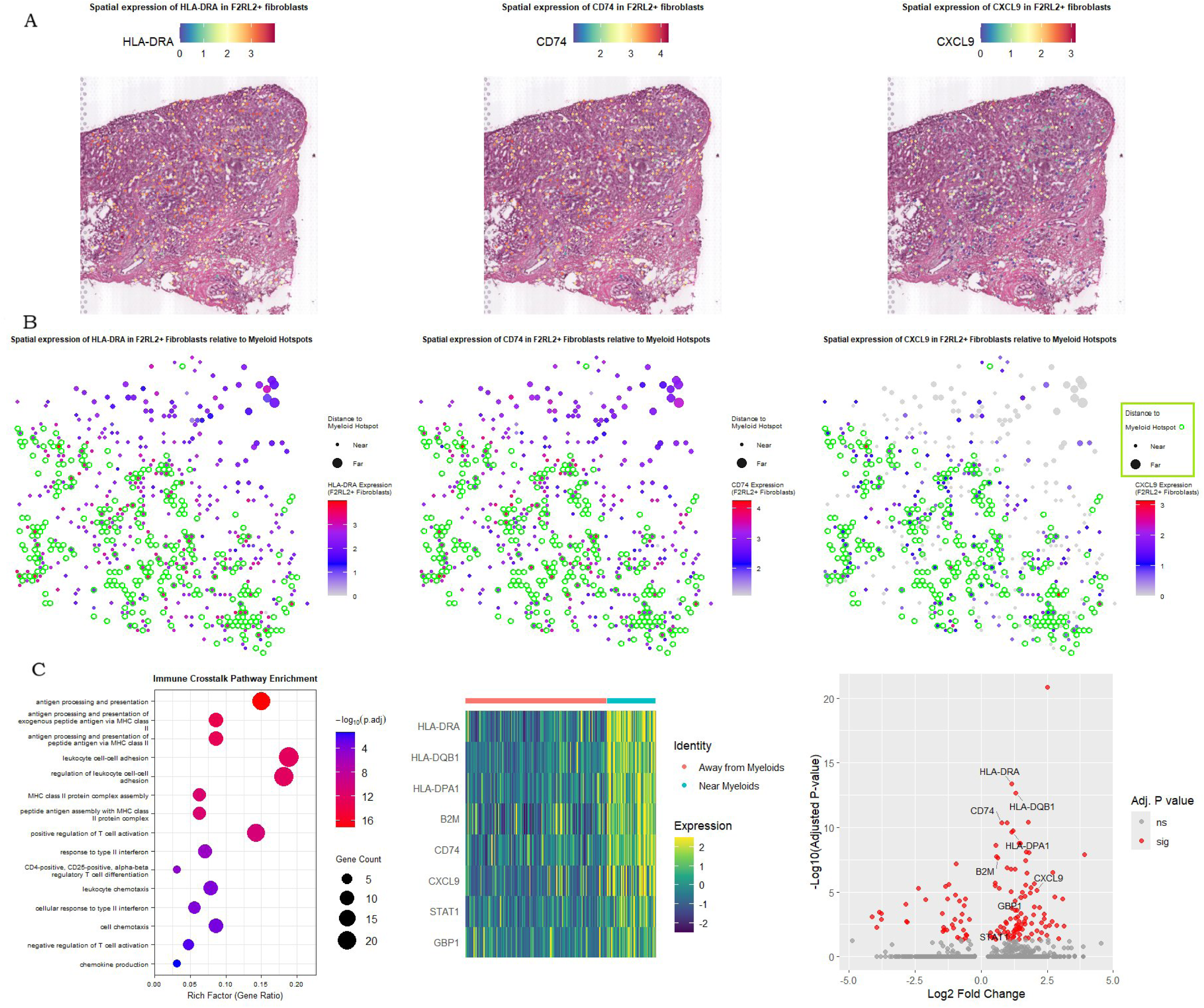
The spatial expression patterns of key immune-related genes in F2RL2+ fibroblasts relative to myeloid hotspots in tissue T2. The figure shows the spatial expression patterns of key genes involved in antigen presentation and T-cell regulation in F2RL2+ fibroblasts relative to myeloid hotspots. (A) Spatial expression of HLA-DRA, CD74, and CXCL9 in F2RL2+ fibroblasts. The color scales indicate the expression levels of these genes. (B) Spatial distribution of HLA-DRA, CD74, and CXCL9 expression in F2RL2+ fibroblasts relative to myeloid hotspots. The scatter plots show the relationship between gene expression and distance from myeloid hotspots. (C) Immune crosstalk pathway enrichment analysis. The heatmap and bar plot summarize the enrichment of pathways related to antigen presentation, T-cell regulation, and leukocyte adhesion in the F2RL2+ fibroblast population. The analysis reveals that F2RL2+ fibroblasts located near myeloid cells show robust upregulation of MHC-II machinery (HLA-DRA, CD74) and interferon-response genes (CXCL9, STAT1), with significant enrichment of pathways involved in antigen presentation via MHC class II, T-cell regulation, and leukocyte adhesion. (hypergeometric testing and BH correction. via clusterProfiler)

In T7, proximal F2RL2+ fibroblasts upregulated MHC-II genes (HLA-DRA, P _adj_ = 2.9e-4; CD74, P _adj_ = 3.3e-7) and adhesion markers (MYL9, P _adj_ = 8.4e-3; ZYX, P _adj_ = 1.9e-2), but chemokine signaling was non-significant, leading to enriched cytokine production (GO:0002361, P _adj_ = 6.1e-3) and interferon response (GO:0034341, P _adj_ = 6.1e-3). By contrast, F2RL2+ spots in T3 and T6 showed no significant immune enrichment despite myeloid proximity (all P _adj_ > 0.05), indicating context-dependent loss of crosstalk. COL6A6+ fibroblasts exhibited minimal immune engagement. (Supplementary Figure 7)

In T2, proximal spots showed trend-level MHC-II upregulation (HLA-DRA, P = 5.9e-6) but no significant pathways after adjustment (P _adj_ > 0.05). (Figure 6 A&B) In T3, T6, and T7, extreme scarcity (≤5 spots per sample) precluded detection of immune activation. MAB21L1+ fibroblasts uniformly lacked significant immune pathway enrichment across tumors (all P _adj_ > 0.05), with insufficient spots for analysis in T6/T7 (≤2 per sample).

These results establish that functional immune activation requires direct myeloid-fibroblast contact (<100μm), and that patient-specific disruption occurs through two mechanisms: biochemical uncoupling (F2RL2 in T3/T6) or cellular depletion (COL6A6). This spatial licensing paradigm explains how proximity-dependent crosstalk collapse contributes to metastatic niche permissiveness.

### 5. Clinical and Therapeutic Relevance

#### 5.1 Genomic and prognostic validation

CNAs analysis of 3,878 primary and 578 metastatic breast cancers revealed distinct alteration patterns (Table 2). Metastases exhibited significantly higher rates of deep deletions for MAB21L1 (9.13% vs. 0.57% in primaries) and COL6A6 (4.76% vs. 0.03% in primaries). F2RL2 also showed increased deep deletions in metastases (2.38% vs. 0.42% in primaries). Amplifications were less frequent but followed similar trends, with MAB21L1 (2.51% vs. 0.82%) and COL6A (1.46% vs. 0.76%) showing higher rates in metastases. These genomic patterns suggest metastatic progression preferentially selects for loss of MAB21L1, F2RL2, and COL6A6, supporting their potential role as metastasis suppressors (Table 2, Supplementary Figure 8).

**Table 2:** Copy Number Alterations (CNA) and Mutation Frequencies in Primary Breast Cancer (PBC) and Breast Cancer (BC) Metastasis.

| Gene | PBC<br>(%CNA)<br>(N:3878) | BC Metastasis<br>(%CNA)<br>(N:578) | CNA: Deep Deletion, N (%) |  | CNA: Amplification, N (%) |  |
| --- | --- | --- | --- | --- | --- | --- |
|  |  |  | PBC<br>(N:3530) | BC Metastasis<br>(N:756) | PBC<br>(N:3530) | BC Metastasis<br>(N:756) |
| **MAB21L1** | 1% | 15% | 20 (0.57%) | 69 (9.13%) | 29 (0.82%) | 19 (2.51%) |
| **F2RL2** | <1% | 5% | 15 (0.42%) | 18 (2.38%) | 8 (0.23%) | 13 (1.72%) |
| **COL6A6** | 1% | 11% | 1 (0.03%) | 36 (4.76%) | 27 (0.76%) | 11 (1.46%) |

BC: Breast Cancer; CNA: Copy Number Alteration; N: Number; PBC: Primary Breast Cancer.

Kaplan-Meier analysis demonstrated significant prognostic associations for three genes in lymph node-negative, systemically untreated breast cancer patients. Patients with high expression of **COL6A6** (n=39 high vs. 18 low; HR = 0.19, 95% CI: 0.06–0.62; log-rank p = 0.0022) exhibited an 81% reduction in relapse risk compared to the low-expression group. Similarly, high expression of F2RL2 (n=36 high vs. 21 low; HR = 0.3, 95% CI: 0.09– 0.94; p = 0.028) was associated with a 70% lower risk of relapse. For MAB21L1, high expression (n=655 high vs. 310 low; HR = 0.66, 95% CI: 0.53–0.83; p = 0.00027) correlated with a 34% risk reduction. All survival curves showed statistically significant separation (log-rank p < 0.05), underscoring the potential role of these genes as biomarkers for favorable prognosis and metastasis suppression in breast cancer (Figure 7).

**Figure 7.**
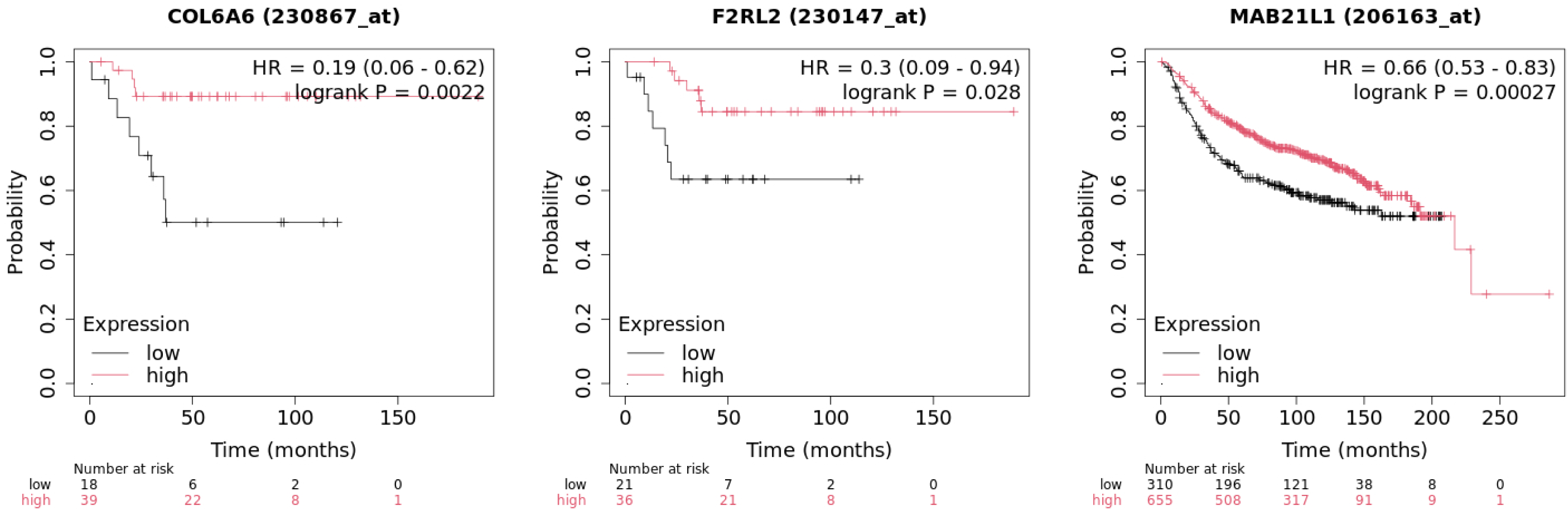
Kaplan-Meier survival analyses demonstrating the prognostic significance of three genes - COL6A6, F2RL2, and MAB21L1 - in lymph node-negative, systemically untreated primary breast cancer patients. (A) High expression of COL6A6 (n=39 high vs. 18 low) was associated with an 81% reduction in relapse risk compared to the low-expression group (HR = 0.19, 95% CI: 0.06–0.62, log-rank p = 0.0022). (B) High expression of F2RL2 (n=36 high vs. 21 low) was associated with a 70% lower risk of relapse (HR = 0.3, 95% CI: 0.09–0.94, log-rank p = 0.028). (C) High expression of MAB21L1 (n=655 high vs. 310 low) correlated with a 34% reduction in relapse risk (HR = 0.66, 95% CI: 0.53–0.83, log-rank p = 0.00027). All three survival curves showed statistically significant separation (log-rank p < 0.05), underscoring the potential of these genes as biomarkers for favorable prognosis and metastasis suppression in lymph node-negative, systemically untreated breast cancer. (Log-rank test (for Kaplan-Meier curve significance, Cox proportional hazards model (for hazard ratios).

TNMplot analysis showed unique expression trends for three candidate genes in breast cancer. COL6A6 showed a robust decrease in expression between normal tissue (median expression = 466) and primary tumor (median = 26; fold change [FC] = 0.12, p = 4.87×10) and an even larger decrease when comparing metastasis (median = 7; FC = 0.44 vs. tumor; p = 4.58×10 ²). MAB21L1 expression also decreased from normal (median = 158) to tumor (median = 16; FC = 0.16, p = 5.23×10), while the decrease to metastasis (median = 15; FC = 0.91 vs. tumor, p = 2.93×10 ¹) was not statistically significant. F2RL2 had contrasting expression with a large increase (median = 480 vs. normal = 132; FC = 3.21; p = 9.51×10 ¹), then was significantly decreased in metastasis (median = 13; FC = 0.03 vs. tumor; p = 1.04×10). These expression profiles propose that COL6A6 and MAB21L1 are metastasis suppressors, while F2RL2 is context-dependent during cancer progression (Table 3, Supplementary Figure 9). Identifying Candidate

**Table 3:** Differential Expression of Candidate Genes in Breast Cancer Progression.

| GENE | FOLD<br>CHANGE | FOLD<br>CHANGE | P-VALUE<br>(NL-T) | P-VALUE<br>(T-METS) | MEDIAN<br>EXPRESSION | MEDIAN<br>EXPRESSION | MEDIAN<br>EXPRESSION |
| --- | --- | --- | --- | --- | --- | --- | --- |

|  | (T/NL) | (METS/T) |  |  | (NL) | (T) | (METS) |
| --- | --- | --- | --- | --- | --- | --- | --- |
| **COL6A6** | 0.12 | 0.44 | $4.87 \times 10^{-56}$ | $4.58 \times 10^{-2}$ | 466 | 26 | 7 |
| **F2RL2** | 3.21 | 0.03 | $9.51 \times 10^{-18}$ | $1.04 \times 10^{-5}$ | 132 | 480 | 13 |
| **MAB21L1** | 0.16 | 0.91 | $5.23 \times 10^{-59}$ | $2.93 \times 10^{-1}$ | 158 | 16 | 15 |

METS: METASTATIC; NL: NORMAL; T: TUMOR.

#### 5.2 Therapeutic candidates for stromal restoration

Our analysis identified candidate drugs that target either the downregulated genes MAB21L1, F2RL2, and COL6A6 in LNMs of breast cancer. For F2RL2, the top candidate drugs were the mRNA expression increasing Valproic Acid and Tretinoin, with Valproic Acid having the additional target of epigenetic modulation through the inhibition of HDAC. Another candidate drug justifiably prioritized was Cyclosporine. This drug is FDA approved for immunosuppression, but it would increase F2RL2 expression and could domain a specific role for F2RL2 support a potential immune-modulatory role in human breast cancer (Table 4).

**Table 4:**
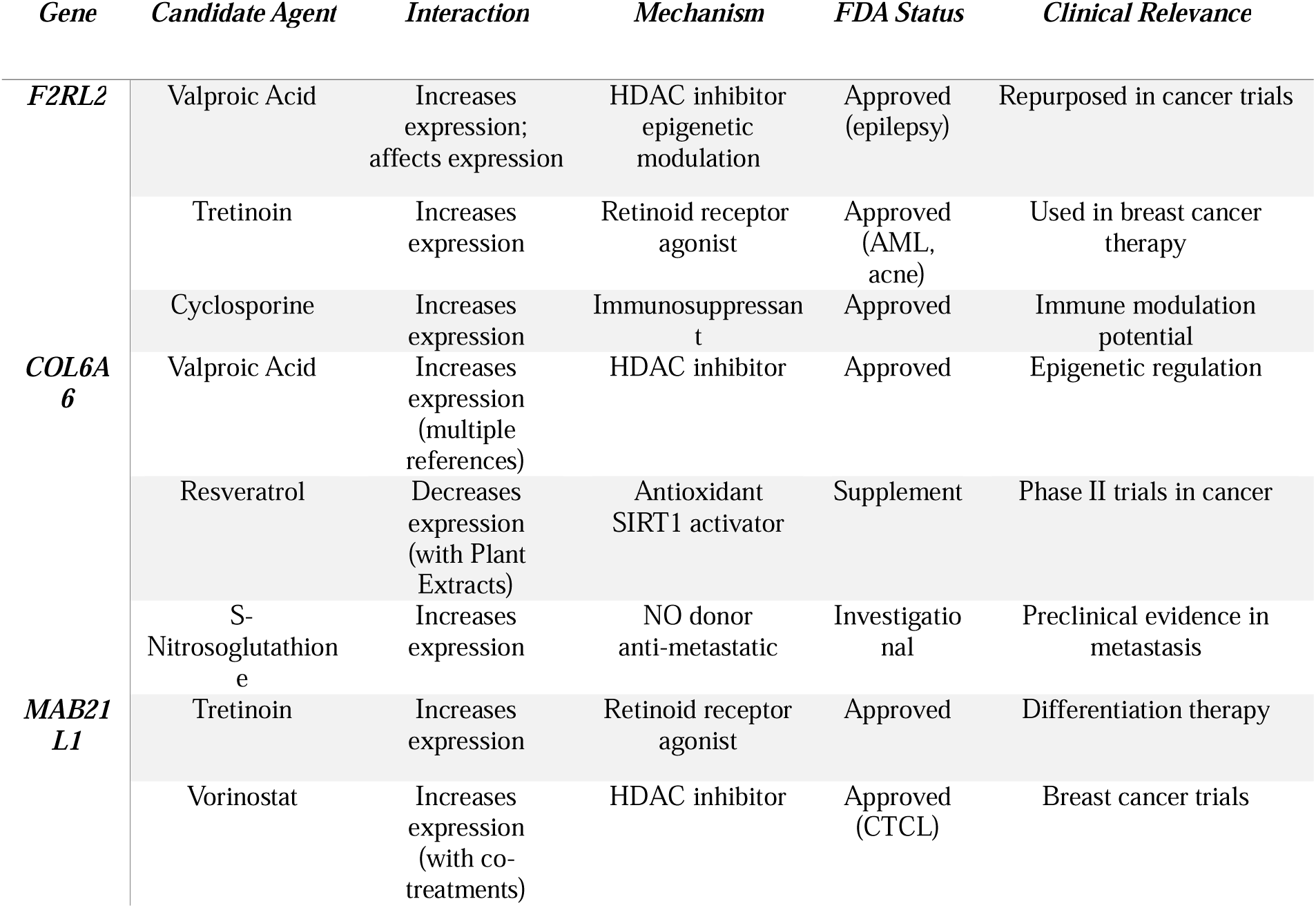
Prioritized Therapeutic Candidates Targeting MAB21L1, F2RL2, and COL6A6.

## Discussion

This multi-omics study systematically identifies conserved molecular drivers of breast cancer lymph node metastasis (LNM) through an unbiased transcriptome-first approach. We discovered MAB21L1, F2RL2, and COL6A6 as consistently downregulated genes in LNMs across independent cohorts—revealing their suppression as a hallmark of metastatic adaptation. Single-cell and spatial resolution demonstrated that these genes define specialized fibroblast subpopulations orchestrating complementary metastasis-constraining mechanisms within primary tumors. F2RL2 fibroblasts spatially coordinate immune-metastasis crosstalk through CXCL14-mediated monocyte recruitment and C1S-initiated complement activation, while balancing TGF-β/BMP signaling to limit epithelial-mesenchymal transition. COL6A6 fibroblasts fortify physical barriers via collagen VI assembly and suppress pro-dissemination pathways, including Wnt signaling. MAB21L1 fibroblasts inhibit stromal-tumor signaling by blocking SMAD-dependent pathways.

These subpopulations collectively establish a multi-layered stromal defense system that physically, immunologically, and biochemically constrains metastatic dissemination. Clinical validation confirms their role as metastasis suppressors, with genomic deletion enrichment in metastases (CNA analysis), stage-specific downregulation (TNMplot), and significant association between high expression and prolonged recurrence-free survival (KM analysis). The erosion of these specialized fibroblast niches in LNMs—marked by subpopulation depletion, ECM dissolution, and loss of immune coordination—creates a permissive microenvironment for metastatic progression. This work defines a paradigm of fibroblast functional diversification in metastasis suppression and nominates candidate therapeutic agents (e.g., Valproic Acid, Tretinoin) for stromal-targeted intervention.

In metastatic niches, we observed a coordinated **stromal collapse** characterized by three pathological shifts: (1) Depletion of metastasis-constraining fibroblast subsets (F2RL2 : 1.9% → 0.1%; COL6A6 : 3.1% → 1.3%); (2) Pathogenic reprogramming of residual fibroblasts toward ECM dissolution (loss of COL6A6/DCN/FBLN5) and aberrant immune modulation (CCL19/21 induction); and (3) Spatial disorganization that disrupts proximity-dependent immune crosstalk. Critically, F2RL2 fibroblasts—which license MHC-II and chemokine expression (e.g., CXCL9) only when positioned <100μm from myeloid cells in primary tumors—lose this immune-activating capacity in LNMs. This erosion of spatial licensing creates permissive niches for metastatic outgrowth, aligning with recent paradigms of microenvironmental coordination in metastatic suppression [9,10].

Clinically, genomic analyses revealed selective pressure for suppressor loss in metastases, with significantly elevated deep deletion frequencies for MAB21L1 (9.13% vs. 0.57% in primaries) and COL6A6 (4.76% vs. 0.03%). In line with previous research suggesting that COL6A6 inhibits cancer cell progression through the JAK signaling pathway and the PI3K-Akt pathway, and is positively associated with immune cell infiltration, our findings indicate that high COL6A6 expression is associated with an 81% reduction in relapse risk (HR=0.19, p=0.0022). This further underscores its role as an early gatekeeper against dissemination. [11–14] MAB21L1 downregulation in lymph node metastases disrupts stromal defense mechanisms and promotes metastatic survival, potentially through loss of its anti-apoptotic function. This is mechanistically supported by findings demonstrating MAB21L1’s role in suppressing the ATR/CHK1/p53 pathway and regulating Bcl-2 family proteins to protect against apoptosis. [15] Paradoxically, F2RL2 exhibited stage-dependent dynamics: upregulation in primary tumors (FC=3.21 vs. normal) followed by profound suppression in metastases (FC=0.03). This suggests context-dependent roles, potentially reflecting its dual function in stromal barrier maintenance versus microenvironmental plasticity [16, 17]

Therapeutically, computational screening prioritized agents capable of restoring suppressive stroma, including epigenetic modulators (Valproic Acid, Vorinostat) and retinoids (Tretinoin) that upregulate the identified genes. While requiring experimental validation, these candidates highlight druggable pathways to reinforce stromal containment. Future studies should functionally validate these fibroblast subpopulations in metastatic restraint using organotypic models and explore whether stromal collapse mechanisms extend to visceral metastases.

In conclusion, our unbiased approach—initially agnostic to cellular mechanisms—converged on specialized fibroblast subpopulations as central executors of molecular programs constraining LN dissemination. The spatial and functional erosion of these fibroblasts enables metastatic progression through multi-layered stromal collapse. Rather than broadly targeting CAFs, selectively restoring these suppressive functions represents a promising therapeutic strategy. Our integrated spatial atlas provides a blueprint for developing microenvironment-focused interventions against breast cancer metastasis.

For clarity, all reported p-values were adjusted using **Benjamini-Hochberg (BH)** unless otherwise specified. Non-parametric tests were prioritized due to non-normal data distributions.

