## Supplementary File for "Spatial and Single-Cell Dissection of Fibroblast Subpopulation Reprogramming Driving Stromal Collapse in Breast Cancer Lymph Node Metastasis"

**Supplementary Table 1**

| **Cell Type** | **Tissue Type** | **MAB21L1_Negative** | **MAB21L1_Positive** | **F2RL2_Negative** | **F2RL2_Positive** | **COL6A6_Negative** | **COL6A6_Positive** |
| --- | --- | --- | --- | --- | --- | --- | --- |
| B cell | LymphNode | 6790 | 2 | 6791 | 1 | 6790 | 2 |
| B cell | Primary | 596 | 0 | 596 | 0 | 596 | 0 |
| Endothelial cell | LymphNode | 2928 | 2 | 2928 | 2 | 2929 | 1 |
| Endothelial cell | Primary | 13720 | 31 | 13742 | 9 | 13742 | 9 |
| Epithelial cell | LymphNode | 53 | 0 | 53 | 0 | 53 | 0 |
| Epithelial cell | Primary | 5633 | 8 | 5638 | 3 | 5636 | 5 |
| Fibroblast | LymphNode | 1308 | 24 | 1331 | 1 | 1315 | 17 |
| Fibroblast | Primary | 3718 | 105 | 3749 | 74 | 3706 | 117 |
| Myeloid cell | LymphNode | 536 | 1 | 536 | 1 | 537 | 0 |
| Myeloid cell | Primary | 4673 | 2 | 4668 | 7 | 4671 | 4 |
| NK cell | LymphNode | 671 | 0 | 670 | 1 | 671 | 0 |
| NK cell | Primary | 363 | 0 | 360 | 3 | 363 | 0 |
| Plasma cell | LymphNode | 148 | 0 | 148 | 0 | 148 | 0 |
| Plasma cell | Primary | 234 | 0 | 234 | 0 | 234 | 0 |
| T cell | LymphNode | 16908 | 1 | 16884 | 25 | 16904 | 5 |
| T cell | Primary | 7034 | 5 | 7016 | 23 | 7034 | 5 |

**Supplementary Table 2:** Differential gene expression (DEG) prevalence across fibroblast subpopulations

| **Subcluster** | **Cell Count** | **MAB21L1 (%)** | **F2RL2 (%)** | **COL6A6 (%)** |
| --- | --- | --- | --- | --- |
| 0 | 828 | 0.367 | 0 | 0 |
| 1 | 672 | 8.30 | 8.59 | 10.1 |
| 2 | 526 | 1.95 | 0.177 | 0 |
| 3 | 482 | 0.639 | 0 | 0 |
| 4 | 404 | 3.12 | 0.284 | 0.284 |
| 5 | 370 | 0.637 | 0 | 0 |
| 6 | 366 | 2.15 | 1.08 | 0 |
| 7 | 356 | 7.09 | 6.38 | 19.1 |
| 8 | 355 | 3.77 | 2.83 | 17.9 |
| 9 | 247 | 2.11 | 1.05 | 2.11 |
| 10 | 187 | 0 | 0 | 0 |
| 11 | 147 | 0 | 0 | 0 |
| 12 | 124 | - | - | - |
| 13 | 56 | - | - | - |
| 14 | 35 | - | - | - |

**Notes:**

- **Cell Count**: Total number of fibroblasts in each subcluster (from fibroblasts_all$seurat_clusters).
- **DEG Prevalence (%)**: Percentage of cells expressing each gene within the subcluster (from deg_prevalence tibble).
- **Missing DEG Data**: Subclusters **12, 13, 14** were not included in the DEG prevalence analysis.
- **0% Values**: Indicates no detectable expression in the subcluster.

*Supplementary Table 3:*

| **Subcluster** | **Gene Expression** | **Key Pathways & Functions** | **Biological Role Summary** |
| --- | --- | --- | --- |
| **1** | High expression of **all 3 genes** | **1. Protein Homeostasis & Stress Response** - Protein refolding, chaperone-mediated folding - Response to heat/unfolded protein - PERK-mediated UPR (Integrated Stress Response)  **2. mRNA Regulation** - mRNA catabolism (deadenylation, poly(A) tail shortening) - Primary miRNA processing  **3. Inflammation & Immunity** - Chronic inflammatory response - Regulation of NOD2/NLR signaling  **4. Cell Differentiation** - Negative regulation of stem cell/adipocyte differentiation - Odontoblast/cardiac neural crest differentiation | **Protein quality control + immune/stress signaling** Maintains proteostatic resilience under cellular stress |
| **7** | Highest **COL6A6**; High **F2RL2/MAB21L1** | **1. ECM Organization** - ECM assembly/disassembly - Collagen fibril/basement membrane organization  **2. TGF-β/BMP Signaling** - TGF-β receptor superfamily pathway - BMP signaling - Cellular response to TGF-β/BMP  **3. Wound Healing & Remodeling** - Regulation of angiogenesis - Wound healing - Bone development/remodeling  **4. Immune Modulation** - Complement activation (classical/alternative) - Humoral immune response  **5. Growth Factor Signaling** - PI3K/Akt pathway - VEGF signaling - Regulation of ERK cascade | **ECM remodeling + TGF-β-driven tissue organization** Aligned with COL6A6's core ECM functions |
| **8** | Moderate expression; High **COL6A6** | **1. Mitochondrial Energy Metabolism** - Oxidative phosphorylation - Electron transport chain - ATP synthesis  **2. Inflammatory Response** - Respiratory burst in inflammation - Leukocyte migration/aggregation  **3. Cellular Respiration** - Aerobic respiration - Proton transmembrane transport  **4. ROS Dynamics** - Response to ROS - ROS metabolic processes | **Metabolic support + inflammatory modulation** Supports metabolic adaptation and immune crosstalk (e.g., with myeloid cells) |

*Supplementary Table 4:* Confirmation of Tumor Microenvironment Implications

| **Key Implication Point** | **Support from Enrichment Analysis** |
| --- | --- |
| **Subcluster 7: ECM Dysregulation → Desmoplasia** | *"Subcluster 7... dominant enrichment for extracellular matrix (ECM) organization (collagen fibril assembly, basement membrane formation)"* → **Confirms ECM dysregulation as a driver of desmoplastic stroma.** |
| **Subcluster 8: Metabolic Reprogramming → Nutrient Support** | *"Subcluster 8... linked to mitochondrial energy metabolism (oxidative phosphorylation, ATP synthesis)"* → **Validates metabolic reprogramming for tumor bioenergetics.** |
| **Subcluster 1: Chronic Inflammation → Immune Evasion** | *"Subcluster 1... enriched for immune/stress regulation (chronic inflammatory response, NOD2 signaling)"* → **Aligns with inflammation-driven immune suppression.** |
| **LN Metastasis: Loss of Stromal Programs** | *"Collective downregulation in LNMs signifies loss of critical stromal programs—ECM integrity (Sc7), stress resilience (Sc1), metabolic/immune coordination (Sc7/8)"* → **Directly explains how pathway suppression enables metastasis.** |
| **Reduced ECM Barriers** | *"Loss of ECM integrity (Subcluster 7)"* → Facilitates cancer cell invasion. |
| **Attenuated Immune Signals** | *"Loss of immune coordination (Subclusters 7 & 8)"* → Enables evasion of surveillance. |
| **Metabolic Adaptation** | *Non-redundant expression underscores specialized contributions* → Explains niche-specific survival in LNMs. |

*Supplementary Table 5*: *Key Fibroblast Subtype Markers in Primary Tumors vs. Lymph Node Metastases*

| **Subtype** | **Gene** | **Direction (LNM vs Primary)** | **Magnitude of Change** | **p_val** | **p_val_adj** | **Functional Role** |
| --- | --- | --- | --- | --- | --- | --- |
| **F2RL2+ Fibroblasts** (Descriptive comparison) |  |  |  |  |  |  |
|  | CCL19 | Up | Δ = -4.75 | NA | NA | Lymphoid chemokine |
|  | CCL21 | Up | Δ = -4.50 | NA | NA | Lymphoid chemokine |
|  | CCL2 | Up | Δ = -4.04 | NA | NA | Monocyte recruitment |
|  | MGP | Down | Δ = 3.24 | NA | NA | ECM mineralization |
|  | COL3A1 | Down | Δ = 2.95 | NA | NA | Collagen structure |
|  | DCN | Down | Δ = 2.62 | NA | NA | ECM organization |
|  | POSTN | Down | Δ = 2.40 | NA | NA | Matrix remodeling |
| **COL6A6+ Fibroblasts** (padj < 0.05) |  |  |  |  |  |  |
|  | SLC26A7 | Up | log₂FC = 7.50 | 1.19e-07 | 4.36e-03 | Ion transport |
|  | ADAMTSL2 | Up | log₂FC = 6.98 | 1.19e-07 | 4.36e-03 | ECM remodeling |
|  | LRRN3 | Up | log₂FC = 6.78 | 2.77e-09 | 1.01e-04 | Neuronal adhesion |
|  | COL6A6 | Down | log₂FC = -14.58 | <1e-100 | <1e-100 | Collagen network |
|  | DPT | Down | log₂FC = -3.65 | 3.47e-13 | 1.27e-08 | ECM assembly |
|  | FBLN5 | Down | log₂FC = -3.49 | 5.16e-07 | 1.89e-02 | Elastic fiber formation |
|  | CFD | Down | log₂FC = -3.69 | 2.38e-08 | 8.72e-04 | Complement system |
| **MAB21L1+ Fibroblasts** (Top markers) |  |  |  |  |  |  |
|  | SLC26A7 | Down | log₂FC = -6.54 | 2.42e-05 | 0.884 | Ion transport |
|  | DES | Down | log₂FC = -6.38 | 3.28e-04 | 1.000 | Myofibroblast marker |
|  | ACTG2 | Down | log₂FC = -6.29 | 3.06e-04 | 1.000 | Actin cytoskeleton |
|  | MIR99AHG | Up | log₂FC = 5.52 | 7.30e-05 | 1.000 | Non-coding RNA |
|  | MUCL1 | Up | log₂FC = 5.48 | 1.58e-04 | 1.000 | Epithelial marker |

**Table Notes:**

1. **Change Representation**:
   - F2RL2+: Δ = Primary_Avg - LN_SingleCell (positive = higher in primary)
   - COL6A6+/MAB21L1+: log₂FC = log2(LNM/Primary) (negative = higher in primary)

*Supplementary Table 6:* **Moran's I Test Results**

| **Table** | **Gene** | **Moran_I** | **Expected** | **Variance** | **Z_score** | **p_value** | **Notes** |
| --- | --- | --- | --- | --- | --- | --- | --- |
| T2 | MAB21L1 | 0.0282 | -3e-04 | 1e-04 | 2.6030 | 0.0046 |  |
| T2 | COL6A6 | 0.0203 | -3e-04 | 1e-04 | 1.8834 | 0.0298 |  |
| T2 | F2RL2 | 0.0080 | -3e-04 | 1e-04 | 0.7535 | 0.2256 |  |
| T3 | MAB21L1 | -0.0003 | -4e-04 | 0e+00 | 0.9965 | 0.1595 |  |
| T3 | F2RL2 | 0.0063 | -4e-04 | 1e-04 | 0.5650 | 0.2860 |  |
| T6 | F2RL2 | 0.0179 | -3e-04 | 1e-04 | 1.7665 | 0.0387 |  |
| T7 | F2RL2 | 0.0547 | -6e-04 | 2e-04 | 3.8126 | 0.0001 |  |
| T7 | COL6A6 | -0.0120 | -6e-04 | 2e-04 | -0.8016 | 0.7886 |  |
| T7 | MAB21L1 | -0.0007 | -6e-04 | 0e+00 | -0.8887 | 0.8129 |  |

*Supplementary Table 7:* **Permutation Test Results and Distances** *(Median distances of DEG+ spots to niches with significance from permutation tests)*

| **Tissue** | **Gene** | **# Positive Spots** | **Niche** | **Median Distance (µm)** | **Significant Proximity (BH-adjusted p<0.05)** |
| --- | --- | --- | --- | --- | --- |
| T3 | F2RL2 | 60 | Myeloid | 200.0 | None |
|  |  |  | Tumor | 200.0 | None |
|  |  |  | T_cell | 1715.4 | None |
|  | COL6A6 | 11 | Myeloid | 529.7 | None |
|  |  |  | Tumor | 0.0 | **Tumor (p=0)** |
|  |  |  | T_cell | 2464.4 | None |
|  | MAB21L1 | 13 | Myeloid | 0.0 | **Myeloid (p=0)** |
|  |  |  | Tumor | 200.0 | None |
|  |  |  | T_cell | 1119.4 | **T_cell (p=0)** |
| T2 | F2RL2 | 379 | Myeloid | 200.2 | **Myeloid (p=0)** |
|  |  |  | Tumor | 200.2 | None |
|  |  |  | T_cell | 531.5 | None |
|  | COL6A6 | 85 | Myeloid | 200.0 | **Myeloid (p=0)** |
|  |  |  | Tumor | 200.3 | None |
|  |  |  | T_cell | 529.7 | **T_cell (p=0)** |
|  | MAB21L1 | 56 | Myeloid | 201.2 | None |
|  |  |  | Tumor | 200.0 | None |
|  |  |  | T_cell | 531.6 | None |
| T6 | F2RL2 | 39 | Myeloid | 200.0 | **Myeloid (p=0)** |
|  |  |  | Tumor | 200.0 | None |
|  |  |  | T_cell | 1060.9 | **T_cell (p=0)** |
|  | COL6A6 | 5 | Myeloid | 200.0 | **Myeloid (p=0)** |
|  |  |  | Tumor | 200.0 | None |
|  |  |  | T_cell | **1443.8** | None |
|  | MAB21L1 | 9 | Myeloid | 200.0 | **Myeloid (p=0)** |
|  |  |  | Tumor | 200.2 | None |
|  |  |  | T_cell | 724.2 | **T_cell (p=0)** |
| T7 | F2RL2 | 239 | Myeloid | 200.2 | **Myeloid (p=0)** |
|  |  |  | Tumor | 200.0 | None |
|  |  |  | T_cell | 721.8 | **T_cell (p=0)** |
|  | COL6A6 | 21 | Myeloid | 200.0 | **Myeloid (p=0)** |
|  |  |  | Tumor | 347.2 | None |
|  |  |  | T_cell | 401.7 | **T_cell (p=0)** |
|  | MAB21L1 | 10 | Myeloid | 200.1 | **Myeloid (p=0)** |
|  |  |  | Tumor | 200.7 | None |
|  |  |  | T_cell | 465.9 | **T_cell (p=0)** |

*Supplementary Table 8:* **Correlation with Markers** *(Spearman correlation of DEGs with immune/epithelial markers; only significant correlations shown)*

| **Tissue** | **Gene** | **Marker** | **Rho** | **p-value** | **Direction** | **Cell Type Association** |
| --- | --- | --- | --- | --- | --- | --- |
| **T3** | F2RL2 | – | – | – | – | No significant correlations |
|  | COL6A6 | EPCAM | 0.058 | 0.003 | Positive | Epithelial |
|  |  | KRT19 | 0.045 | 0.021 | Positive | Epithelial |
|  | MAB21L1 | CD68 | 0.047 | 0.015 | Positive | Myeloid |
|  |  | NKG7 | 0.046 | 0.016 | Positive | NK cell |
|  |  | CD3D | 0.040 | 0.037 | Positive | T cell |
| **T2** | F2RL2 | LYZ | 0.099 | 2.8e-08 | Positive | Myeloid |
|  |  | CD68 | 0.048 | 0.007 | Positive | Myeloid |
|  |  | CD3D | 0.089 | 5.2e-07 | Positive | T cell |
|  |  | MRC1 | 0.036 | 0.045 | Positive | Myeloid (M2) |
|  |  | NKG7 | 0.036 | 0.043 | Positive | NK cell |
|  |  | KRT19 | -0.036 | 0.042 | Negative | Epithelial |
|  | COL6A6 | NKG7 | 0.045 | 0.011 | Positive | NK cell |
|  |  | EPCAM | -0.048 | 0.007 | Negative | Epithelial |
|  | MAB21L1 | CD14 | 0.049 | 0.006 | Positive | Myeloid |
|  |  | LYZ | -0.056 | 0.002 | Negative | Myeloid |
| **T6** | F2RL2 | LYZ | 0.044 | 0.009 | Positive | Myeloid |
|  |  | CD3D | 0.041 | 0.014 | Positive | T cell |
|  | COL6A6 | – | – | – | – | No significant correlations |
|  | MAB21L1 | CD3D | 0.051 | 0.002 | Positive | T cell |
| **T7** | F2RL2 | MRC1 | 0.115 | 9.9e-07 | Positive | Myeloid (M2) |
|  |  | LYZ | 0.084 | 3.5e-04 | Positive | Myeloid |
|  |  | CD68 | 0.072 | 0.002 | Positive | Myeloid |
|  |  | CD14 | 0.053 | 0.026 | Positive | Myeloid |
|  |  | CD3D | 0.062 | 0.008 | Positive | T cell |
|  |  | KRT19 | -0.111 | 2.2e-06 | Negative | Epithelial |
|  |  | EPCAM | -0.048 | 0.041 | Negative | Epithelial |
|  | COL6A6 | KRT19 | -0.081 | 6.4e-04 | Negative | Epithelial |
|  |  | EPCAM | -0.066 | 0.005 | Negative | Epithelial |
|  | MAB21L1 | KRT19 | -0.052 | 0.026 | Negative | Epithelial |
|  |  | EPCAM | -0.049 | 0.037 | Negative | Epithelial |

*Supplementary Table 9:* ***Comparison with Random Spots*** *(Proximity of DEG+ spots to myeloid hotspots vs. random spots)*

| **Tissue** | **Gene** | **DEG+ Spots Near Myeloid (%)** | **Random Spots Near Myeloid (%)** | **Absolute Difference (%)** | **p-value** |
| --- | --- | --- | --- | --- | --- |
| **T3** | F2RL2 | 8.3 | 3.3 | 5.0 | 0 |
|  | COL6A6 | 9.1 | 0.03 | 9.1 | 0.001 |
|  | **MAB21L1** | **15.4** | **0.01** | **15.4** | **0** |
| **T2** | F2RL2 | 12.9 | 10.0 | 2.9 | 0 |
|  | COL6A6 | 10.6 | 7.1 | 3.5 | 0.001 |
|  | **MAB21L1** | **5.4** | **3.6** | **1.8** | **0.001** |
| **T6** | F2RL2 | 15.4 | 10.2 | 5.1 | 0 |
|  | COL6A6 | 20.0 | 0.02 | 20.0 | 0.001 |
|  | **MAB21L1** | **11.1** | **0.0** | **11.1** | **0** |
| **T7** | F2RL2 | 13.0 | 9.6 | 3.4 | 0 |
|  | COL6A6 | 4.8 | 9.5 | -4.8 | 1 |
|  | **MAB21L1** | **20.0** | **0.03** | **20.0** | **0.001** |

*Supplementary Figures*

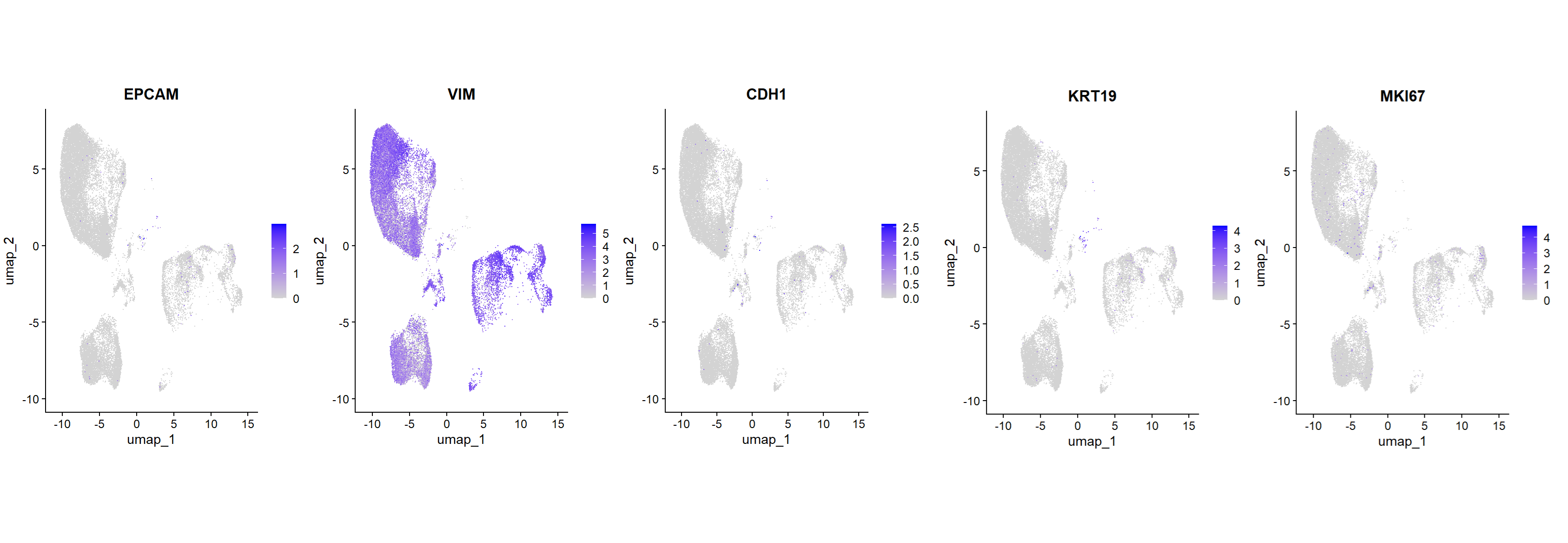

***Supplementary Figure1*** This figure presents feature plots profiling the spatial expression patterns of key markers in lymph node metastases (LNMs): EPCAM: Marks epithelial cells, showing a heterogeneous distribution across the LNM. VIM: Vimentin, a mesenchymal marker, exhibits pan-tissue expression, indicating widespread stromal activation and potential epithelial-mesenchymal plasticity. CDH1: E-cadherin, an epithelial marker, shows a more focused expression pattern within the LNM. KRT19: Keratin 19, another epithelial marker, displays a similar localized expression profile. MKI67: A proliferation marker, is concentrated in specific epithelial compartments, suggesting active proliferation with stromal involvement. These spatial expression patterns reveal three key insights: Widespread vimentin (VIM) expression indicates robust stromal activation and potential epithelial-mesenchymal plasticity across the LNM. Proliferation (MKI67) is concentrated in distinct epithelial regions, with stromal involvement. Epithelial markers (EPCAM, CDH1, KRT19) exhibit more heterogeneous and localized expression patterns within the LNM. Together, these findings provide a comprehensive spatial profile of the cellular composition and functional states within lymph node metastases.

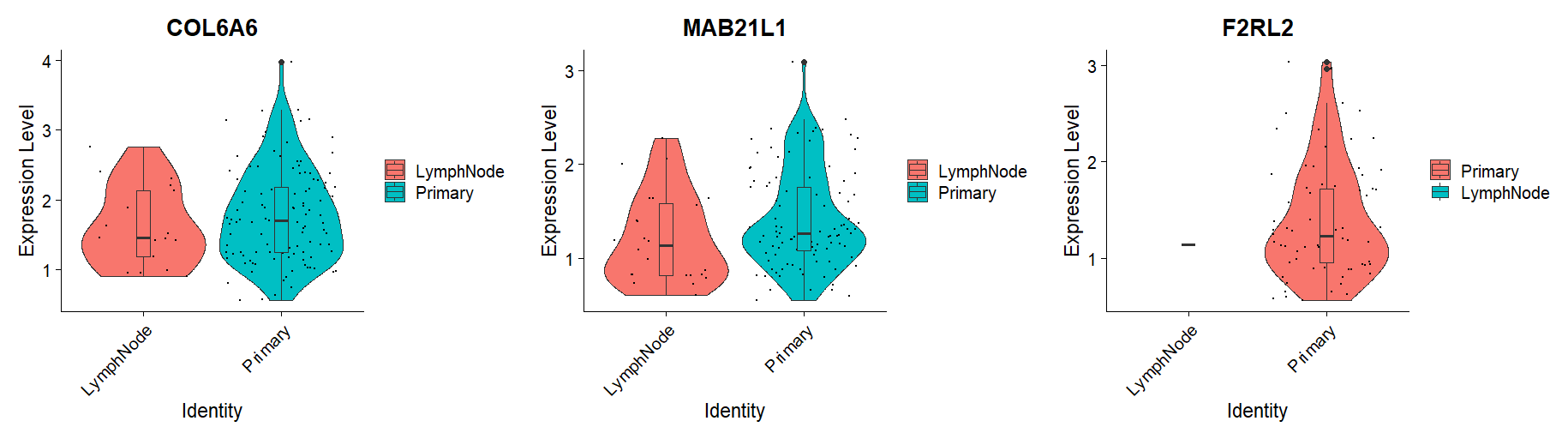

*Supplementary Figure 2. The three metastasis-associated genes (MAB21L1, F2RL2, COL6A6) exhibited significantly higher expression in fibroblast cells of primary tumors compared to lymph node metastases (LNMs). No co-expression was detected among these genes.*

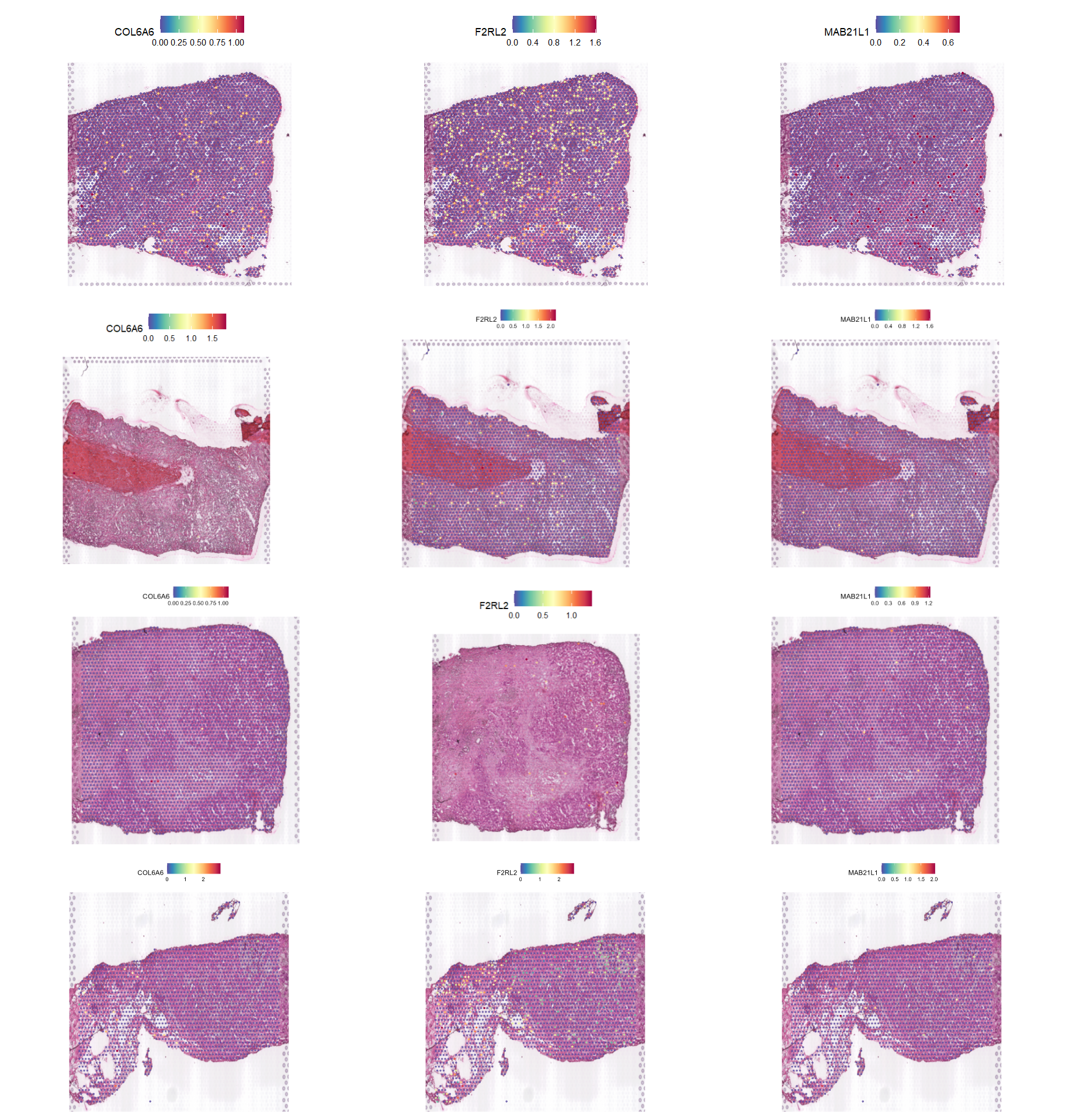

***Supplementary Figure 3:*** *Spatial expression patterns of three metastasis-associated genes (MAB21L1, F2RL2, COL6A6) across four primary breast tumors (T2, T3, T6, T7), analyzed using spatial transcriptomics. Expression values (ranging from 0.0 to 1.6) are displayed for each gene in distinct tumor regions, representing normalized transcript counts or activity scores. The analysis included 3,153 spots in T2, 2,681 in T3, 3,510 in T6, and 1,793 in T7, revealing heterogeneous gene-specific distributions within the tumor microenvironment. Numerical scales reflect relative expression levels, with higher values indicating increased gene activity.*

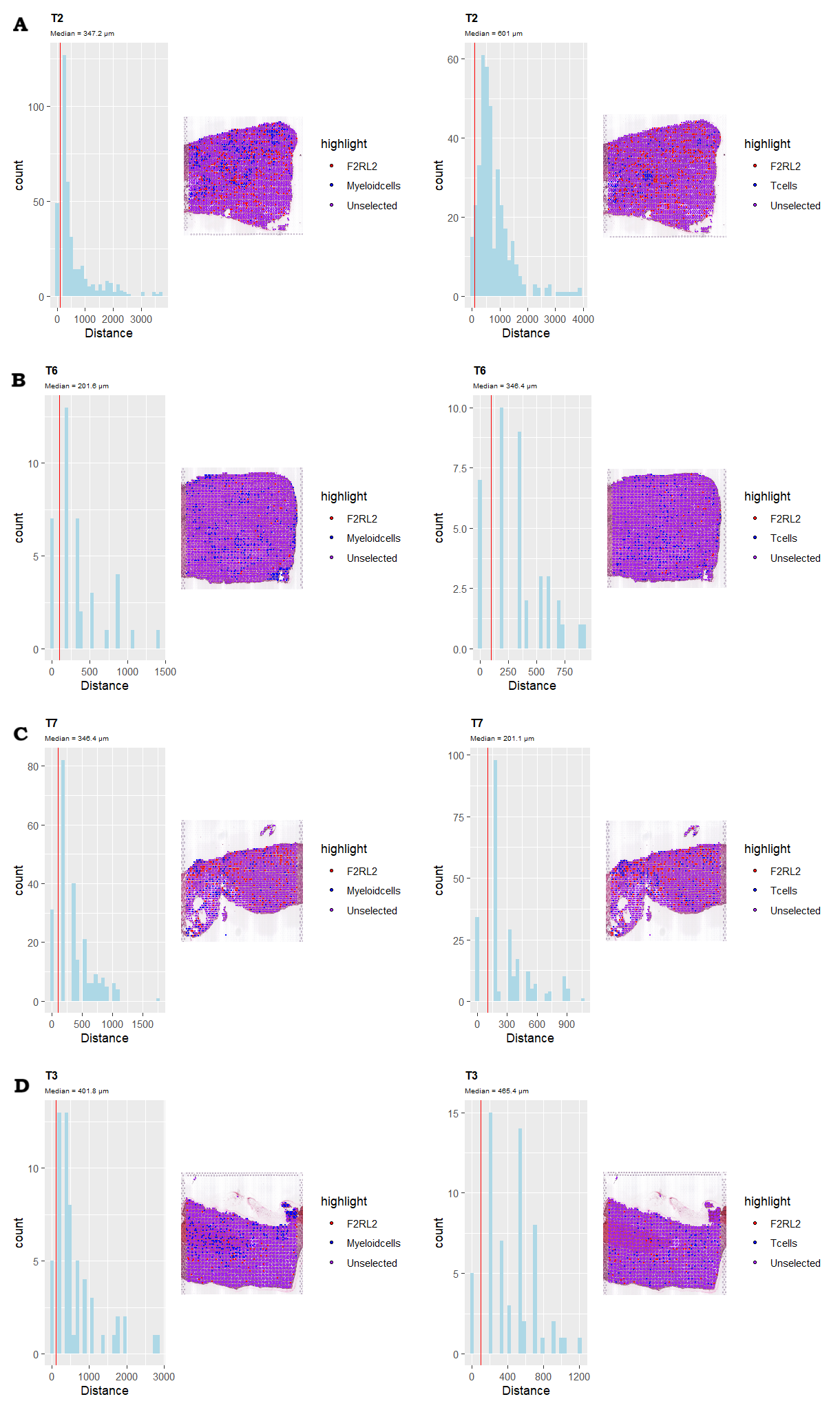

***Supplemantary Figure 4Spatial analysis of F2RL2+ fibroblast proximity to immune niches in sample T2****, T3, T6, and T7. This composite figure compares distances from F2RL2+ fibroblasts to two immune cell populations. Left panels show histograms of minimum distances to myeloid hotspots (top 10% LYZ+ spots) and T-cell hotspots (top 10% CD3D+ spots), with red lines indicating 100 µm thresholds and medians reported. Right panels display spatial maps where myeloid hotspots and ) T-cell hotspots (both blue) are plotted relative to F2RL2+ fibroblasts (red), with purple indicating co-localization.. Distance calculations used k-nearest neighbors (k=5) based on spatial coordinates.*

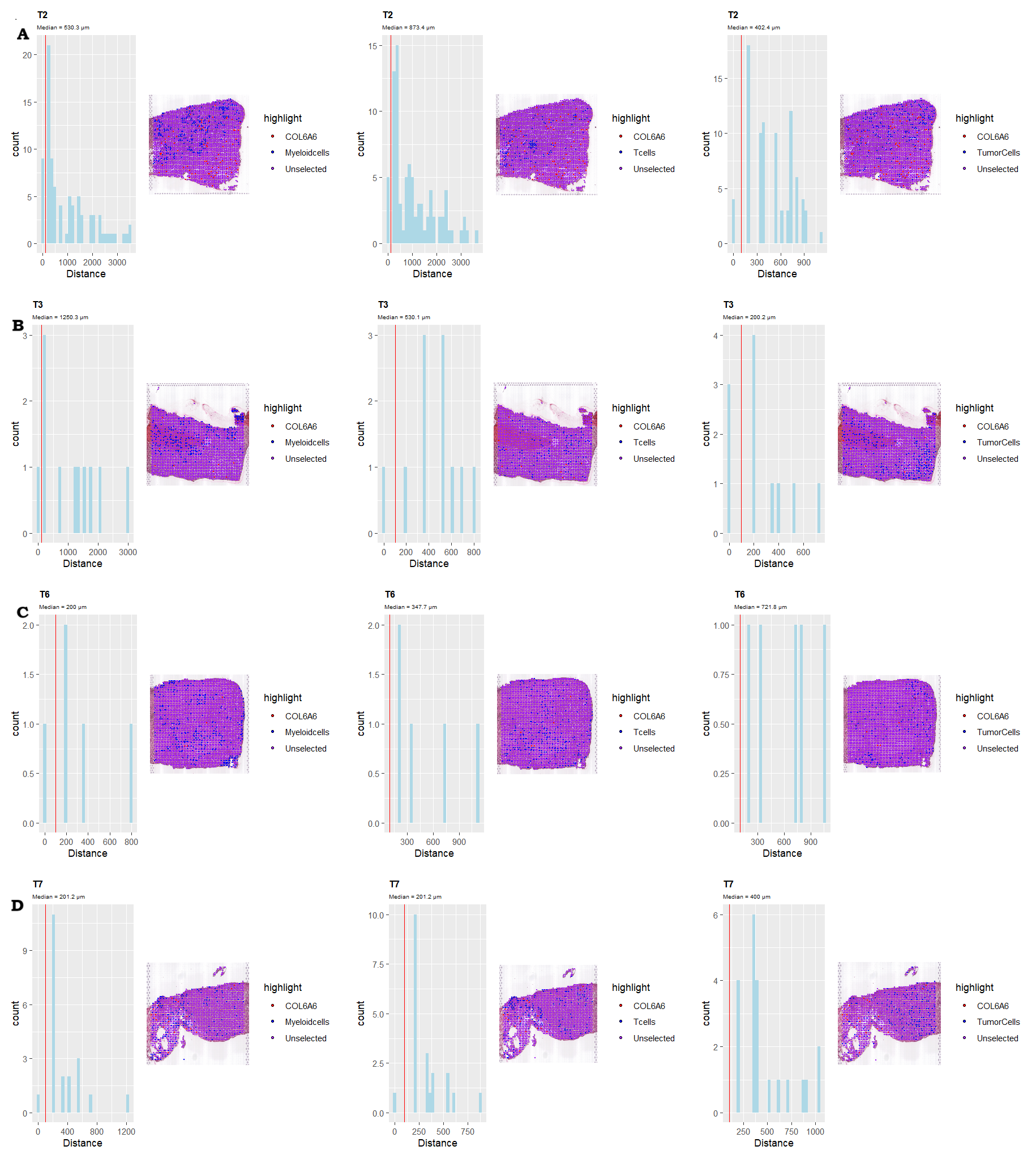

*Supplementary Figure 6*

***Supplemantary Figure 5. Spatial proximity of COL6A6+ fibroblasts to immune/tumor niches in sample T2*** *Each panel pair analyzes distances between COL6A6+ fibroblasts and target cell populations: myeloid hotspots (top 10% LYZ+), T-cell hotspots (top 10% CD3D+), and tumor hotspots (top 10% EPCAM+). Left panels show histograms of minimum distances to each target with 100 µm thresholds (red lines) and reported medians. Right panels display spatial maps where COL6A6+ fibroblasts (red) are plotted relative to target hotspots (blue), with purple indicating co-localization. All distances calculated using k=5 nearest neighbors.*

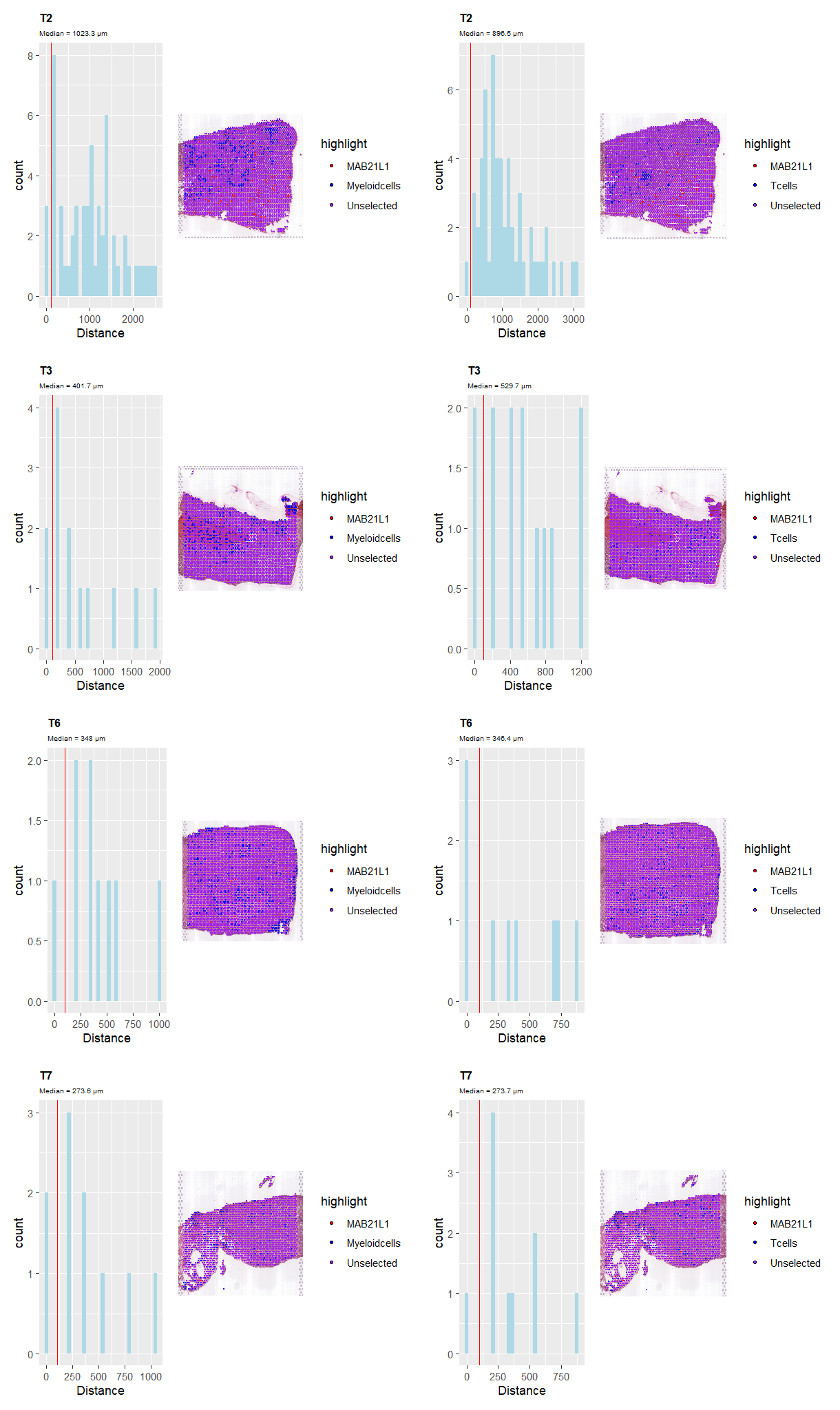

***Supplemantary Figure 4Spatial analysis of MAB21L1+ fibroblast proximity to immune niches in sample T2****, T3, T6, and T7. This composite figure compares distances from MAB21L1+ fibroblasts to two immune cell populations. Left panels show histograms of minimum distances to myeloid hotspots (top 10% LYZ+ spots) and T-cell hotspots (top 10% CD3D+ spots), with red lines indicating 100 µm thresholds and medians reported. Right panels display spatial maps where myeloid hotspots and ) T-cell hotspots (both blue) are plotted relative to MAB21L1+ fibroblasts (red), with purple indicating co-localization.. Distance calculations used k-nearest neighbors (k=5) based on spatial coordinates.*

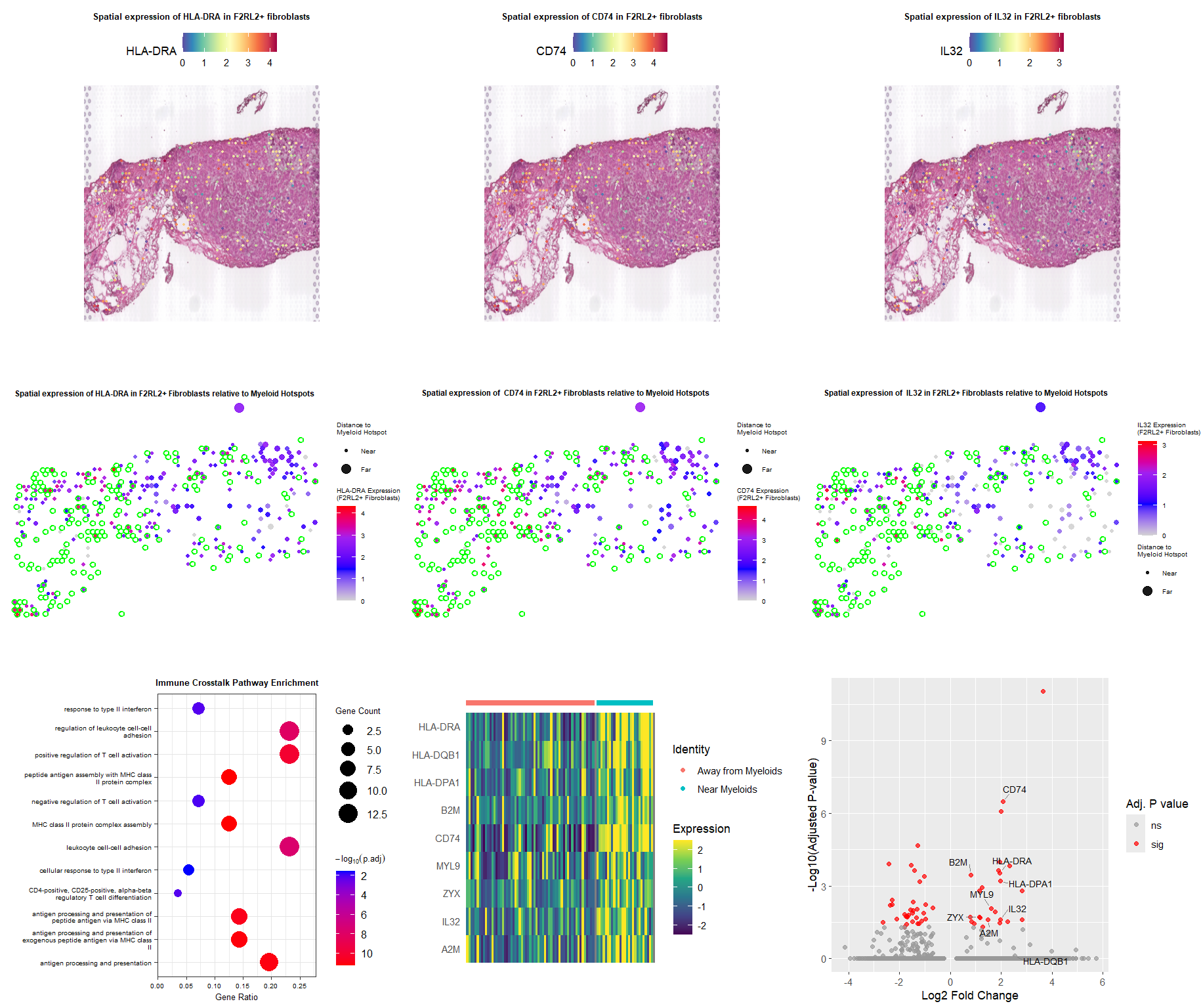

***Supplementary Figure 7.*** ***Context-dependent immune crosstalk in F2RL2+ fibroblasts*** *(A)* ***Spatial expression of MHC-II and adhesion genes in proximal F2RL2+ fibroblasts.***
*Expression patterns of HLA-DRA, CD74 (MHC-II machinery), MYL9, and ZYX (adhesion markers) in F2RL2+ fibroblasts. Color scales denote expression levels. (B)* ***Spatial coordination with myeloid hotspots.*** *Scatter plots depict expression of HLA-DRA, CD74, MYL9, and ZYX relative to distance from myeloid hotspots, highlighting significant upregulation in proximal fibroblasts. (C)* ***Enriched immune pathways and tissue-specific divergence****. Bar plot summarizes pathway enrichment analysis, revealing significant upregulation of cytokine production (GO:0002361, P<sub>adj</sub> = 6.1e-3) and interferon response (GO:0034341, P<sub>adj</sub> = 6.1e-3) in T7. Note: Chemokine signaling pathways were non-significant. In contrast, F2RL2+ fibroblasts in T3/T6 showed no immune enrichment despite myeloid proximity (P<sub>adj</sub> > 0.05), and COL6A6+ fibroblasts exhibited minimal immune engagement.*

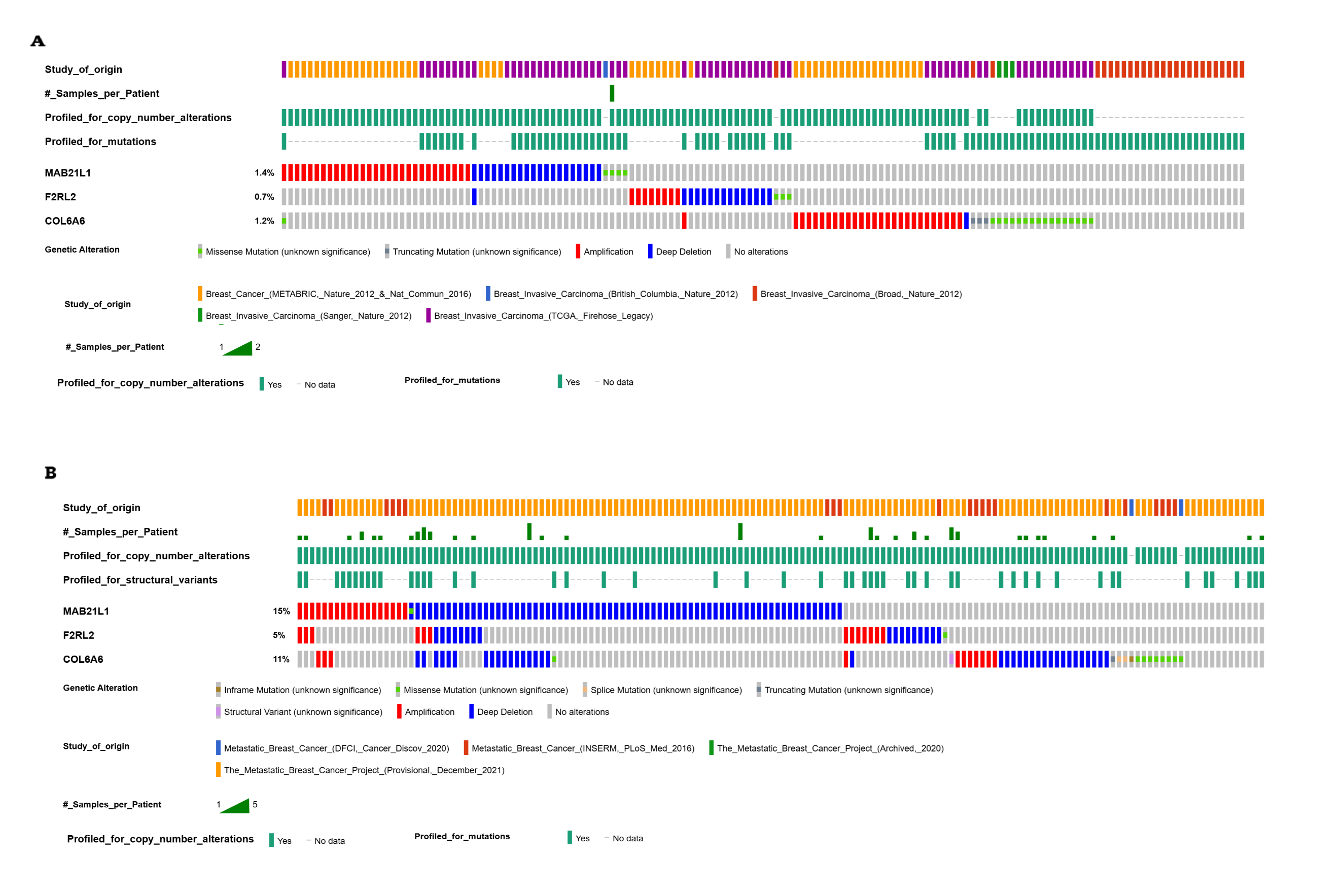

**Supplementary Figure 8.** **Copy number alterations in primary and metastatic breast cancers.** **(A)** Primary breast invasive carcinoma (n= 3,878) showing alterations in MAB21L1, F2RL2, and COL6A6 (altered in 5% of samples). **(B)** Metastatic breast cancer (n = 578) exhibiting significantly higher alteration rates (40% of samples), including deep deletions in MAB21L1 (9.13% vs. 0.57% in primaries), COL6A6 (4.76% vs. 0.03%), and F2RL2 (2.38% vs. 0.42%). Amplifications were less frequent but followed similar trends (MAB21L1: 2.51% vs. 0.82%; COL6A6: 1.46% vs. 0.76%). Data suggests selective loss of these genes in metastases, implicating potential metastasis-suppressor roles.

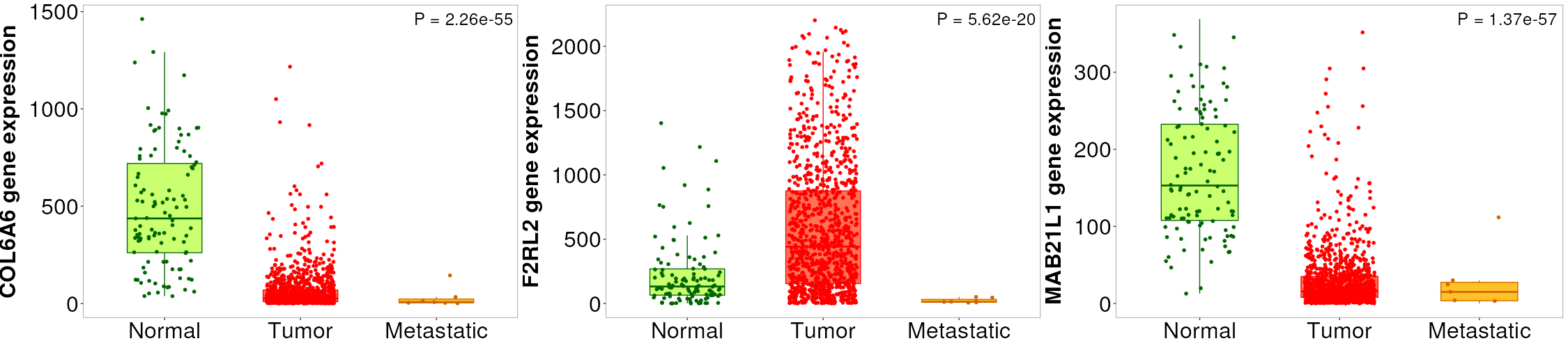

***Supplementary Figure 9: Supplementary Figure 9. TNMplot analysis of gene expression in breast cancer progression.*** *COL6A6 expression ↓ in tumor (****P = 4.87×10⁻⁵⁶****) vs. normal and ↓ further in metastasis (****P* = 4.58×10⁻²****) vs. tumor.
F2RL2 expression ↑ in tumor (****P = 9.51×10⁻¹⁸****) vs. normal but ↓ in metastasis (****P = 1.04×10⁻⁵****) vs. tumor. MAB21L1 expression ↓ in tumor (****P = 5.23×10⁻⁵⁹****) vs. normal; change in metastasis (†P = 0.293) was not significant.*
